# Quantitative and sensitive sequencing of somatic mutations induced by a maize transposon

**DOI:** 10.1101/2025.01.22.634239

**Authors:** Justin Scherer, Michael Hinczewski, Brad Nelms

## Abstract

Cells accumulate mutations throughout development, contributing to cancer, aging, and evolution. Quantitative data on the abundance of *de novo* mutations within plants or animals are limited, as new mutations are often rare within a tissue and fall below the limits of current sequencing depths and error rates. Here, we show that mutations induced by the maize Mutator (Mu) transposon can be reliably quantified down to a detection limit of 1 part in 16,000. We measured the abundance of millions of *de novo* Mu insertions across four tissue types. Within a tissue, the distribution of *de novo* Mu allele frequencies was highly reproducible between plants, showing that, despite the stochastic nature of mutation, repeated statistical patterns of mutation abundance emerge. In contrast, there were significant differences in the allele frequency distribution between tissues. At the extremes, root was dominated by a small number of highly abundant *de novo* insertions, while endosperm was characterized by thousands of insertions at low allele frequencies. Finally, we used the measured pollen allele frequencies to reinterpret a classic genetic experiment, showing that evidence for late Mu activity in pollen are better explained by cell division statistics. These results provide insight into the complexity of mutation accumulation in multicellular organisms and a system to interrogate the factors that shape mutation abundance.

**Significance:** New mutations provide the raw material for evolution and contribute to cancer, aging, and genetic diseases. It has been challenging to follow the origin and spread of new mutations because they can be exceptionally rare and difficult to detect. By focusing on a class of mutation that can be detected more readily – Mu transposon insertions – we followed the abundance of new mutations in multiple maize tissues. We find that the Mu has broad activity across tissues, but with significant tissue-specific differences in how abundant individual new mutations become. Most mutations were below the detection limit available for other classes of mutation. These results provide a glimpse into the complexity of mutation within multicellular organisms.

## INTRODUCTION

Multicellular organisms accumulate mutations throughout development, producing genetic heterogeneity within and between tissues. With the increased sensitivity to detect *de novo* mutations through sequencing, it has become clear that genetic mosaicism is ubiquitous even in healthy individuals^1–8^: over 1,000 single-base substitutions are present per adult human fibroblast^1^ and megabase-sized structural variants can be observed in 30% of healthy human neurons^2^. In plants, low frequency mutations can be transmitted to the next generation^3^, and preexisting (somatic) mutations contribute to variation between plants regenerated in tissue culture^4^.

To interpret and predict the effect of *de novo* mutations, it is critical to understand what influences their abundance and spread within the organism. This is challenging for both biological and technical reasons. Biologically, mutation accumulation is complex and depends on processes that impact the initial mutation rate (e.g. mutagenic exposure, DNA repair) as well as the spread of mutations once they arise (e.g. cell division, selection, cell death). While there have been theoretical advances in understanding how these factors interact to shape mutation abundance^9–15^, there is need for quantitative, empirical data to constrain and inform the theory.

This is where the technical challenge comes in: new mutations can span many orders of magnitude in abundance, down to 1 per cell, pushing the limits of current sequencing depths and error rates. To date, genome-wide studies on *de novo* mutations in plants and animals have reported detection limits around 1-5%^2–8^ (**Fig. S1A**), which cover only the most abundant mutations. Targeted sequencing has helped bridge this gap^16–20^, with an inverse relationship between genomic coverage and sensitivity to detect rare mutations (**Fig. S1A**). However, targeted sequencing still suffers from a limited dynamic range, as more abundant mutations are unlikely to occur within a narrow genomic region.

Mutations caused by transposable elements (TEs) play important roles in evolution, contributing to genome-size evolution^21^, alleles selected during crop domestication^22^, and the origin of new genes through exon shuffling^23^. Unlike other classes of mutation, TE insertions introduce defined sequences into the genome that can be targeted by PCR^24^. The potential of this is significant: by selectively amplifying only genome sequences containing the TE, *de novo* insertions can be identified without sequencing through an overwhelming number of wild-type copies at the same location (**Fig. S1B**). This shares advantages of targeted mutation sequencing without needing to focus on a predefined genomic region.

Here, we evaluate the maize Mutator (Mu) transposon as a quantitative model of *de novo* mutation accumulation in multicellular tissues. Mu has long been a valuable model in maize genetics^25–28^ because of its high forward mutation rate, availability of Mu-active and inactive genetic stocks, and ease of identifying Mu insertion locations by sequencing^24,29^. Mu is a class I (DNA) transposon that predominately transposes duplicatively, i.e. transposing to new locations without loss of the donor element^30^; this apparent ‘copy-and-paste’ behavior is thought to be caused by DNA repair pathways that restore the original sequence after transposition^25,26^. Mu transposes into unlinked sites, with no preference to insert near its site of origin^30^.

We find that Mu sequencing can accurately measure the absolute allele frequencies of *de novo* Mu insertions within complex tissues, with a sensitivity, dynamic range, and error rate that are orders of magnitude better than currently possible for single-base substitutions. We then measured the allele frequency distribution for *de novo* Mu insertions in leaf, root, pollen and endosperm. Mu had broad activity in all four tissues, with no evidence for a preference of late insertion in pollen. These results provide a rich dataset with which to test and refine theoretical models of mutation accumulation in multicellular organisms, and highlight the importance of tissue organization in shaping the abundance of *de novo* mutations during development.

## RESULTS

### Sensitive identification of *de novo* TE insertion sites

To identify Mu TE insertions, we established a sequencing assay based on MuSeq^24^, which has been widely used to map Mu insertions in maize genetic stocks^24,31,32^. MuSeq applies nested PCR to specifically amplify and sequence DNA fragments that span the TE-genome boundary (**Fig. S2**). We optimized MuSeq for quantifying the abundance of rare, *de novo* TE insertions that are heterogeneous within a tissue sample. Two key changes were implemented in ‘MuSeq2’; first, we introduced molecule counting by incorporating unique molecular identifiers^33^ (UMIs) during an initial adapter ligation step (**Fig. S2**). This makes it possible to identify and remove PCR duplicates, improving quantitative accuracy. Second, we limited the amplification of non-Mu products using suppression PCR, providing more specific TE amplification with fewer PCR cycles (**Fig. S3**).

To test MuSeq2, we first applied it to seedling leaves from Mu-active and inactive maize lines. Samples were sequenced to a mean of 1.7 and 2.8 million TE-spanning molecules per Mu-active and inactive plant, respectively. For the inactive plants, all Mu elements are expected to map to a fixed set of genomic locations, representing historical TE insertions. Indeed, 99.8% of molecules from Mu-inactive samples mapped to only 29 locations (**Table S1**). In contrast, Mu-active plants had Mu elements mapping to a wide range of new genomic sites (**Fig. S4A,B**), with a mean of 184,432 insertion sites detected per leaf. These can be confirmed as *bona fide* Mu insertions because (i) the TE border was consistently sequenced along with the genomic region (**Fig. S4C**) and (ii) 123,312 sites were sequenced out of both directions of the TE, including the 9 bp target site duplication that is characteristic of Mu insertions^26^. To estimate the error rate of MuSeq2, we leveraged the fact that Mu-inactive lines have a negligible rate of new Mu transposition, providing a genetic control for no transposon activity. Assuming that all molecules mapping outside the 29 historical locations were false positives, the error rate of MuSeq2 is 0.11 ± 0.04 falsely identified insertions per diploid cell (mean ± standard error; N = 3 Mu-inactive plants). This corresponds to 2.7 × 10^−11^ false positive insertions per bp, two orders of magnitude lower than the most accurate duplex methods to measure single-base substitutions^34^.

### *De novo* and inherited Mu insertions across matched tissues

We next applied MuSeq2 to leaf, pollen, endosperm, and root from Mu-active plants. Using sequential tissue isolations and controlled genetic crosses (**Fig. 1A**), we were able to separate *de novo* insertions from inherited ones and further divide the inherited insertions by parent-of-origin. First, the plants were generated from a cross between a Mu-inactive female and Mu-active male; the female parent contributes a defined set of historical insertions (**Fig. 1B**; **Table S1**), and so all other insertion sites in the offspring were either *de novo* or paternally inherited. To distinguish *de novo* from paternally inherited insertions, we used matched tissues with early and well-defined divergence times. The endosperm, which comprises the bulk of maize seed mass, inherits its paternal DNA from a sister sperm cell during double fertilization (**Fig. 1A**, left). Thus, insertions present at high abundance in both endosperm and embryo-derived tissues must be paternally inherited (hereafter: ‘paternal insertions’). Indeed, paternal insertions were well-separated from *de novo* insertions based on their abundance in both endosperm and other tissues from the same plant (**Fig. 1B,C**).

**Figure 1.**
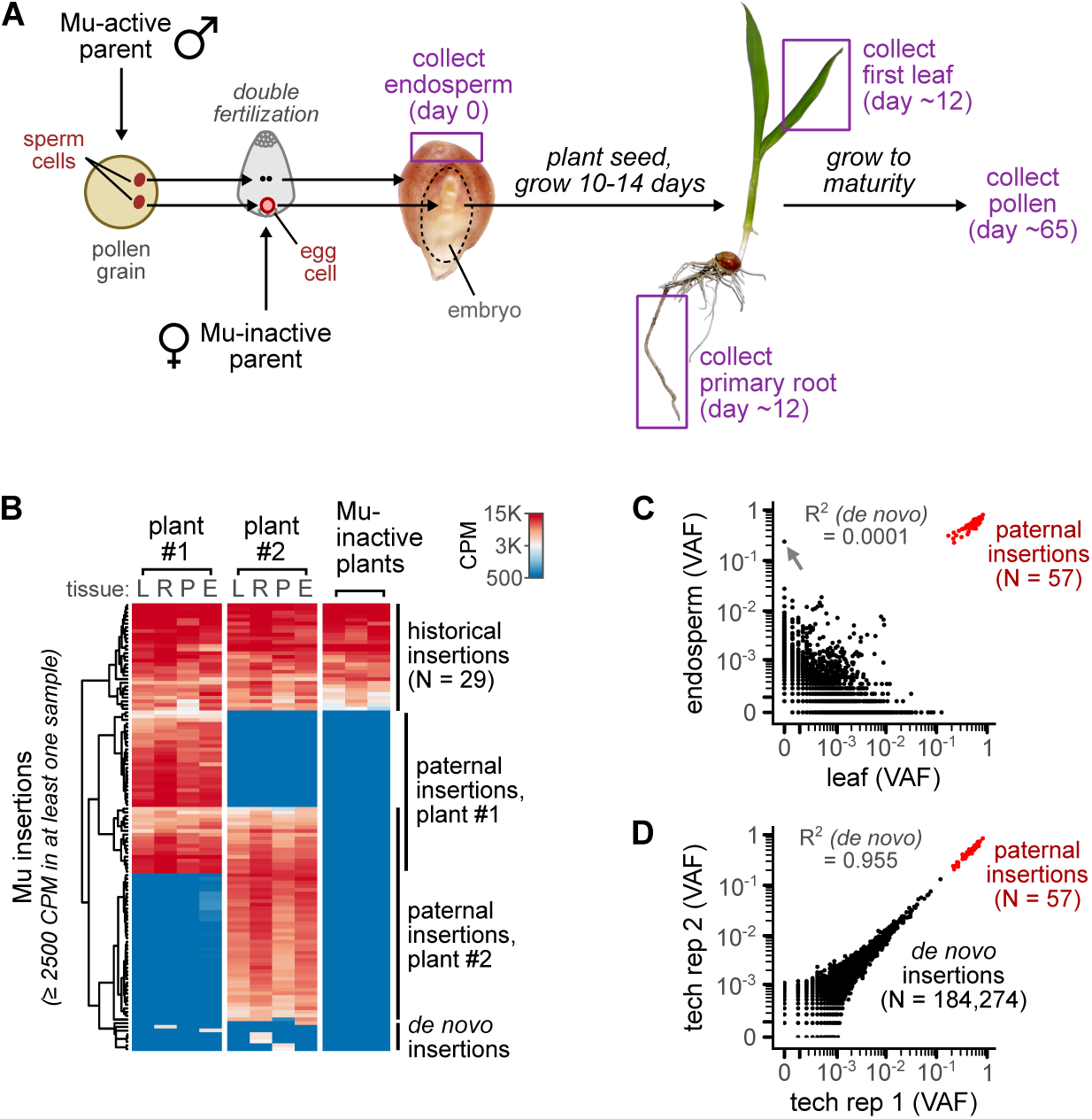
Sensitive and quantitative assessment of *de novo* mutation abundance for an active maize transposon. (**A**) Cartoon of experimental design and tissue collection. Sequential, matched isolations from endosperm and other tissues make it possible to distinguish inherited from *de novo* insertions, because endosperm is derived from a sister sperm cell during double fertilization. (**B**) Heatmap showing the abundance of Mu insertions in matched tissue samples from two siblings as well as control Mu-inactive plants. The Mu-inactive samples were from the family used as the female parent, and represent historical insertions that were maternally inherited. All insertion sites with >= 2500 CPM in at least one of the samples are shown. CPM, counts per million (number of TE-spanning molecules at a given genomic site). (**C**) Allele frequencies of Mu insertions for matched endosperm and leaf from a single plant. Paternal insertions were abundant in both samples, while *de novo* insertions were only abundant in one (e.g. gray arrow). Black dot, *de novo* insertion; red dot, paternal insertion; VAF, variant allele frequency. (**D**) Technical replicates for a representative leaf sample.

The initial output of MuSeq2 is the relative abundance of different Mu insertion sites within a sample. To convert relative abundances (UMI counts) to absolute allele frequencies (variant allele frequency; VAF), we normalized the data using the paternal insertions (**Fig. S4A**), which have a known allele frequency within the sample: 0.5 (heterozygous) in leaf, root and pollen and 0.33 in triploid endosperm (this normalization is insensitive to realistic Mu excision rates; **Fig. S5**). The measured allele frequencies were reproducible between technical replicates (independent libraries prepared from the same DNA; **Fig. 1D**), with strong quantitative agreement across 5 orders of magnitude (R^2^ = 0.997 for all insertions; R^2^ = 0.955 for *de novo* insertions). In contrast, there was no correlation in the abundance of *de novo* insertions between matched tissues from the same plant (**Fig. 1C** and **S6**), reflecting their independent and recent origin. In total, MuSeq2 measured TE allele frequencies down to a detection limit of 6.0 ×10^−5^ for the median sample (1 part in 16,569; *SI Appendix*).

### Allele frequencies measured in pollen match paternal inheritance patterns in the offspring

To benchmark the accuracy of the measured allele frequencies, we compared allele frequencies in bulk pollen to paternal transmission patterns in the offspring (**Fig. 2A**). The number of *de novo* Mu insertions per pollen cell can be calculated as the sum of allele frequencies in pollen (Σv*afi)*. We estimate 22.5 ± 3.8 *de novo* insertions per pollen cell in this Mu-active line (median ± standard error). We then genotyped Mu insertions in the parents and offspring of four families generated from a Mu-active male parent crossed with a Mu-inactive female (N = 30 offspring total, 7-8 per family), making it possible to directly follow the paternal transmission of Mu insertions to the offspring (**Fig. S7**). There were 27.8 ± 6.7 new Mu insertions per offspring, in agreement with the estimate from bulk pollen (**Fig. 2B**). Pollen allele frequencies also agreed with offspring transmission patterns for higher order statistics, such as the number of *de novo* Mu insertions shared by any two siblings (**Fig. 2C** and **S7D**; estimated from pollen as the sum of allele frequencies squared, Σ*vaf_i_^2^*).

**Figure 2.**
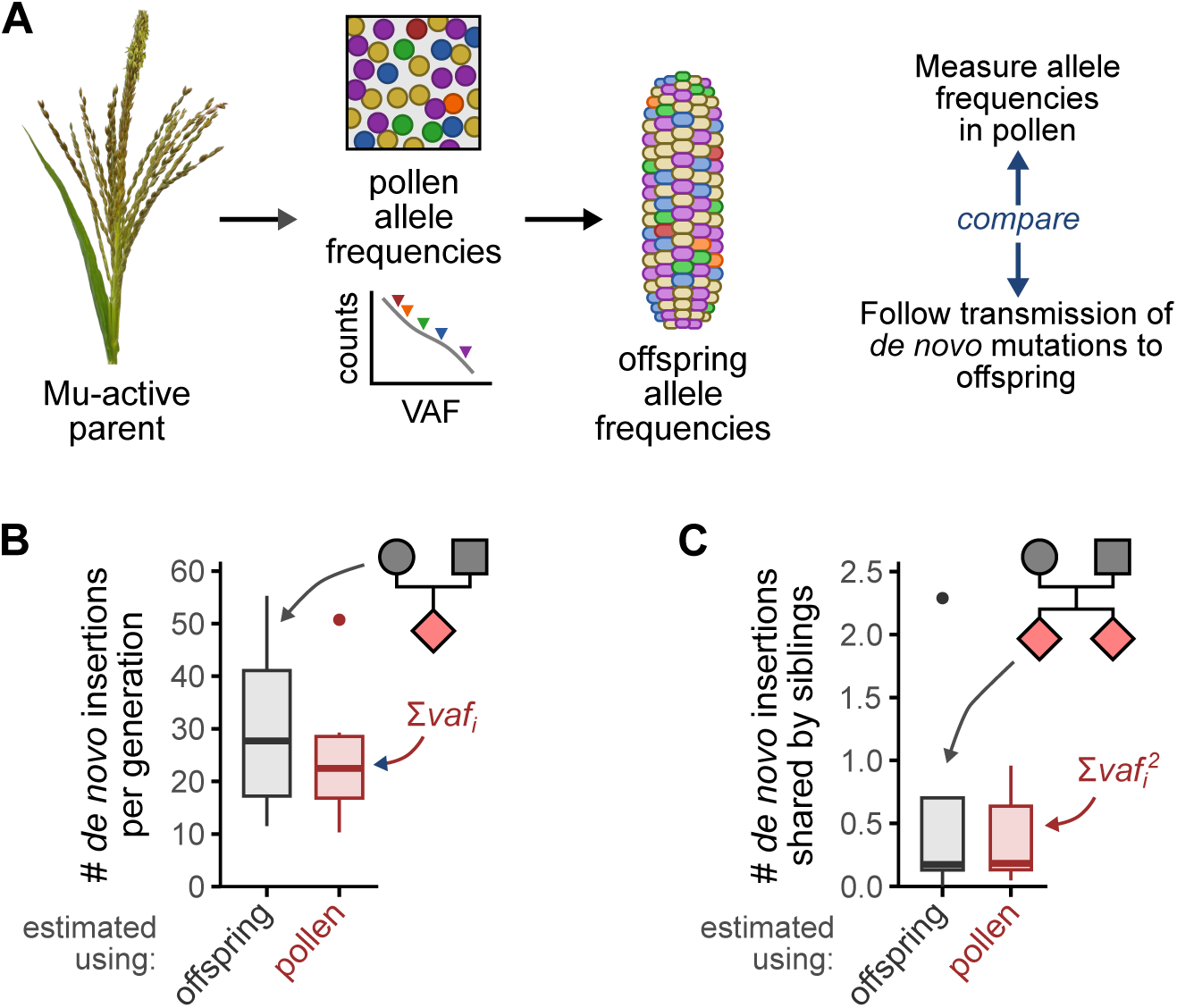
Pollen allele frequencies match paternal inheritance patterns. (**A**) Allele frequencies in pollen should predict allele frequencies in the offspring. (**B**) Number of *de novo* insertions per generation, measured by genotyping parents and offspring (gray; see **Fig. S7A**) or estimated from the pollen allele frequency data (red). The difference between estimates was not significant (p = 0.71, Mann-Whitney U test). (**C**) Number of paternal insertions shared by two siblings, measured by genotyping parents and offspring (gray; see **Fig. S7D**) or estimated from the pollen allele frequency data (red). The difference between estimates was not significant (p = 0.94, Mann-Whitney U test). For panels B and C, N = 4 families with 7-8 offspring each, 9 pollen samples.

To put the Mu transposon activity in context, there are ∼67 single-base substitutions per generation in maize^35^, and similar per base substitution rates have been reported for A*rabidopsis*^36^ and human^37^. Thus, the number of TE insertions in this Mu-active line (20-30 per generation through pollen) is comparable to the background rate of single-base substitutions.

### *De novo* Mu insertions occur at a wide range of allele frequencies

A histogram of Mu allele frequencies for a representative leaf sample is shown in **Fig 3A**. There were 211,097 *de novo* insertions detected in this single leaf, with allele frequencies ranging from 0.28 down to <10^−4^ (the detection limit of the assay). These data suggest that Mu is active throughout development, including insertions that likely arose in the meristem (based on their high abundance^38^; **Fig. 3A**) down to lower frequency insertions that more likely arose in the leaf itself^39^. Mu has a strong preference to insert into and immediately upstream of genes^29^, targeting a reduced portion of the genome. Despite this, we did not observe saturation of the available Mu target sites (**Fig. S8**). The most abundant *de novo* Mu insertions occurred at uncorrelated, independent sites between samples (**Fig. 1C**). Thus, the specific mutations induced by Mutator were stochastic and infrequently repeated between plants. In contrast, the allele frequency distribution was reproducible across its entire range (**Fig. 3B**). Essentially, while the specific set of TE insertions varied widely, a predictable number of insertions were present at any given abundance.

**Figure 3.**
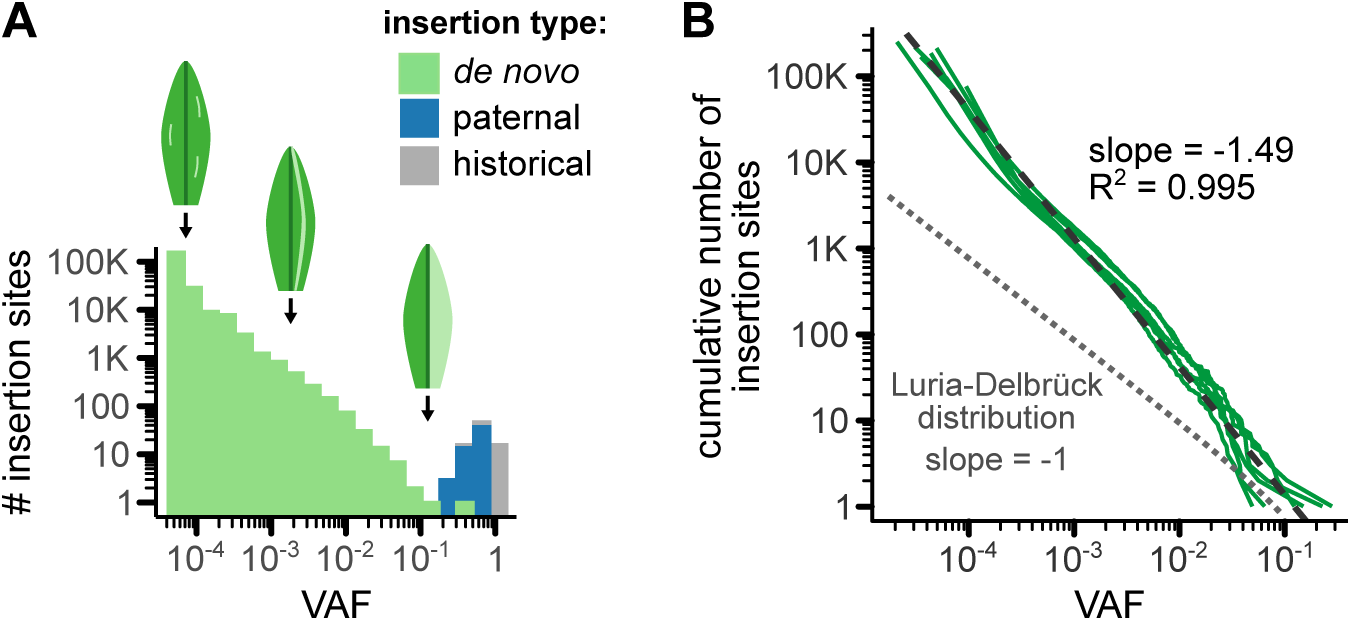
*De novo* Mu insertions occur across a wide range of allele frequencies. (**A**) Histogram of Mu allele frequencies in a representative leaf sample. Colors indicate whether the insertion sites were historical, paternally inherited, or *de novo*. Leaf cartoons show the potential spatial distribution of mutations at selected VAFs, estimated from sector sizes in ref. 39. Insertions above a VAF of 2 × 10^−3^ likely originated in the meristem, based on estimates that 250 meristematic cells form a leaf primordia in maize^38^. VAF, variant allele frequency. (**B**) Cumulative number of Mu insertion sites in individual leaf samples (N = 6). Dashed line, best linear fit to the log-log transformed data; gray dotted line, theoretical expectation for random mutation in an exponentially dividing cell population (Luria-Delbrück distribution).

### Mu insertion activity is much broader than for Mu excisions

Most prior data on somatic Mu activity is based on the excision of Mu elements in the endosperm^25,26,40^, which can be observed by the appearance of revertant purple sectors after Mu excises from an anthocyanin reporter gene (**Fig. S9**). Endosperm excisions produce almost entirely small sectors^40^, suggesting that Mu excision activity is highest later in development^25,26^. The excision rate also varies 1000-fold between tissues, ranging from ∼10% excisions per element in endosperm^40^ down to <10^−4^ excisions per element transmitted through pollen^26^. Compared to excisions, *de novo* Mu insertions were much more broadly distributed across space and time (**Fig. 4A**). There was substantial new insertion activity in every tissue type, despite large divergence in the developmental origins and biology of the selected tissues. Furthermore, *de novo* Mu insertions were observed at a wide range of allele frequencies. While Mu excisions almost never occur above an allele frequency of 0.002^40^, Mu insertions were often observed beyond this limit, even within endosperm (**Fig. 4A** and **S9**). Thus, Mu insertions and excisions behave differently, with new insertions occurring throughout development.

**Figure 4.**
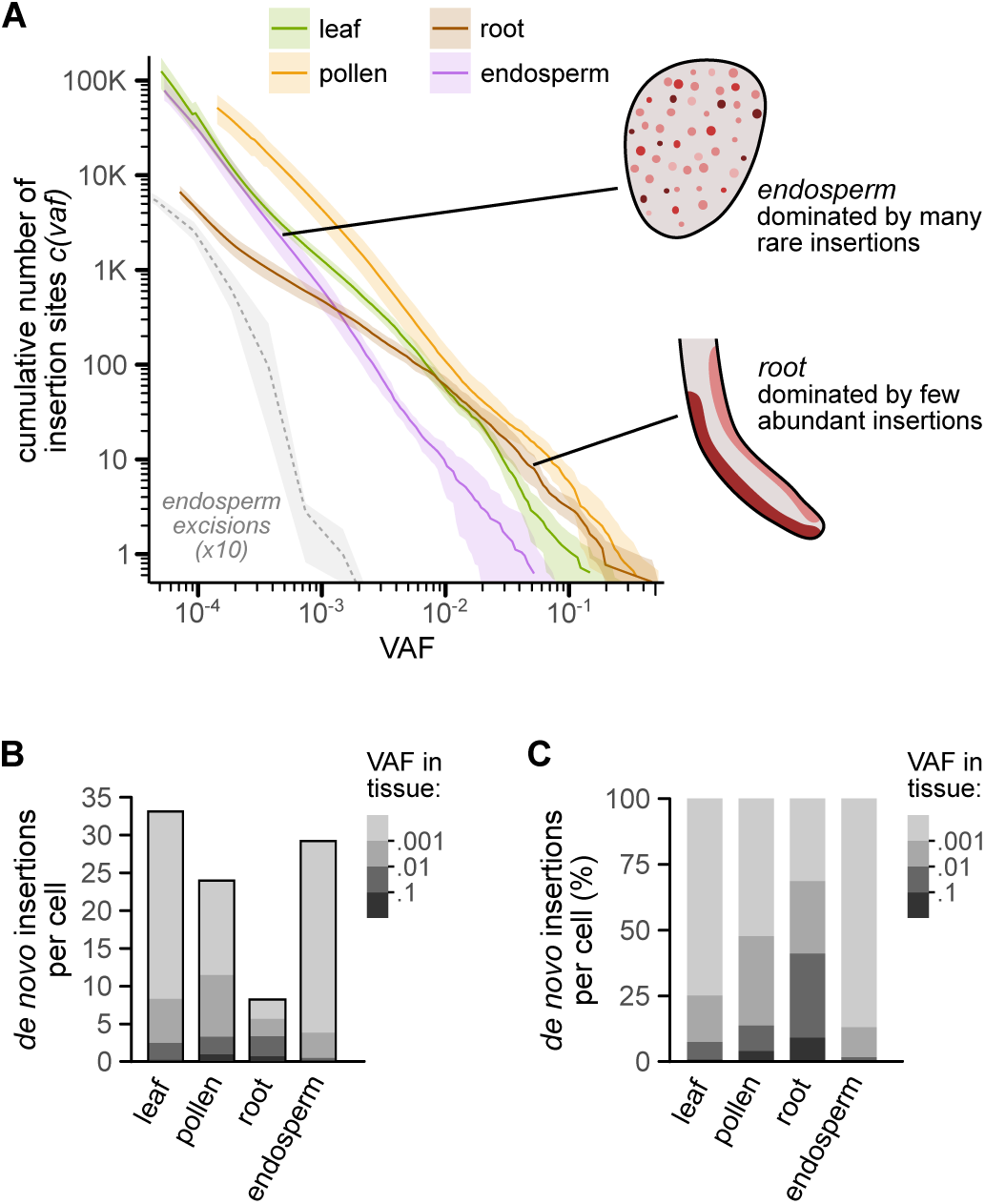
Allele frequency distribution of *de novo* Mu insertions across maize tissue types. (**A**) Cumulative number of *de novo* Mu insertion sites at different allele frequencies. For insertion data (solid lines), curves show the mean and 95% confidence interval (bootstrap test). For endosperm excision data (dotted line), curve shows the reported values; shaded area, 95% confidence interval assuming Poisson counting error. Endosperm excision data is from ref. 40; the reported number of excision events was multiplied by 10 to make the insertion and excision distributions easier to compare. VAF, variant allele frequency. (**B**,**C**) Number and % of *de novo* Mu insertions per cell, calculated from the sum of allele frequencies (Σ*vaf_i_*). These estimates are robust to sequencing depth (**Fig. S10**) and are calculated for cells at the baseline DNA content in each tissue (2C for leaf/root, 3C for endosperm); thus, these are values for a cell that has not endoreduplicated and is in the G1 phase of the cell cycle. Colors indicate the contribution of insertions at different allele frequencies.

### Tissues show distinct allele frequency distributions for *de novo* Mu insertions

While not as dramatic as the divergence between excisions and insertions, there were significant differences in the behavior of *de novo* Mu insertions between tissues (**Fig. 4**). The total number of *de novo* insertions per cell varied by up to 4-fold (**Fig. 4B** and **S10**), ranging from 8.2 in root to 32.8 in leaf. Moreover, each tissue had a reproducible but distinct allele frequency distribution (**Fig. 4A**). Root was most dominated by insertions at high allele frequencies (**Fig. 4B,C**), suggesting that new Mu insertions often formed relatively large sectors in this tissue. At the other extreme, endosperm had a much higher proportion of rare (low VAF) insertions. Thus, there was variation not only in the total number of Mu insertions, but also in how widespread the individual insertions were throughout the tissue.

What might contribute to the observed allele frequencies? Since early studies on bacterial mutation by Luria and Delbrück, many theoretical models of mutation accumulation have been developed. We compared the empirical allele frequency distributions to established theory. Leaf, pollen, and endosperm all closely followed a linear relationship on a log-log plot (a power law distribution; **Fig. 3B** and **S11**). Power-law relationships are well-known in mutation accumulation^14^, as this distribution occurs in an exponentially dividing cell population subjected to a constant rate of neutral mutations (a Luria-Delbrück process). However, the empirical data was a bad fit to the Luria-Delbrück model, because the slopes were far steeper than the theoretical expectation^14^ of −1 (**Fig. 3B**, **S11,** and **S12**). In animals, a common model for mutation accumulation is based on an exponential growth phase early in development, followed by a later, stable-population phase^9,41–44^; however, this model predicts a strong deviation from power law behavior and a shallow slope for much of the range, again a poor fit to the data (**Fig. S13**). Other models of mutation accumulation, including boundary-driven growth^13,14^ (where cells divide preferentially at the edge of an expanding population), linear growth^9^ (such as occurs during asymmetric stem-cell divisions), and the glandular fission model^12^ (developed for solid cancer tumors) also predict sharp deviations from power law behavior. Given the complexity of multicellular development and transposon regulation, it is perhaps unsurprising that established theory cannot explain the data. The availability of quantitative allele frequency data across several orders of magnitude can inform and constrain future theoretical developments to understand mutation accumulation in plants.

### Mu outcross experiments can be explained by cell division statistics

The classic view has been that Mu is most active late in germinal development^25–28^, with activity peaking around the time of meiosis or during pollen maturation. In contrast, our data suggests that Mu insertions occurred throughout development (**Fig. 4A**), with >90% of insertions occurring prior to meiosis (*SI Appendix*). The most direct evidence for Mu activity late in germinal development comes from a study by Robertson^27^, in which he outcrossed Mu-active plants and characterized new mutations in the F1 offspring (**Fig. 5A**). He identified 177 mutant F1 plants that segregated recessive seedling phenotypes; then, through extensive complementation testing, determined that 82.8% of the F1 offspring had unique mutations. The frequent occurrence of unique mutations among the offspring led to the idea that Mu must be most active late in development^25–28^.

**Figure 5.**
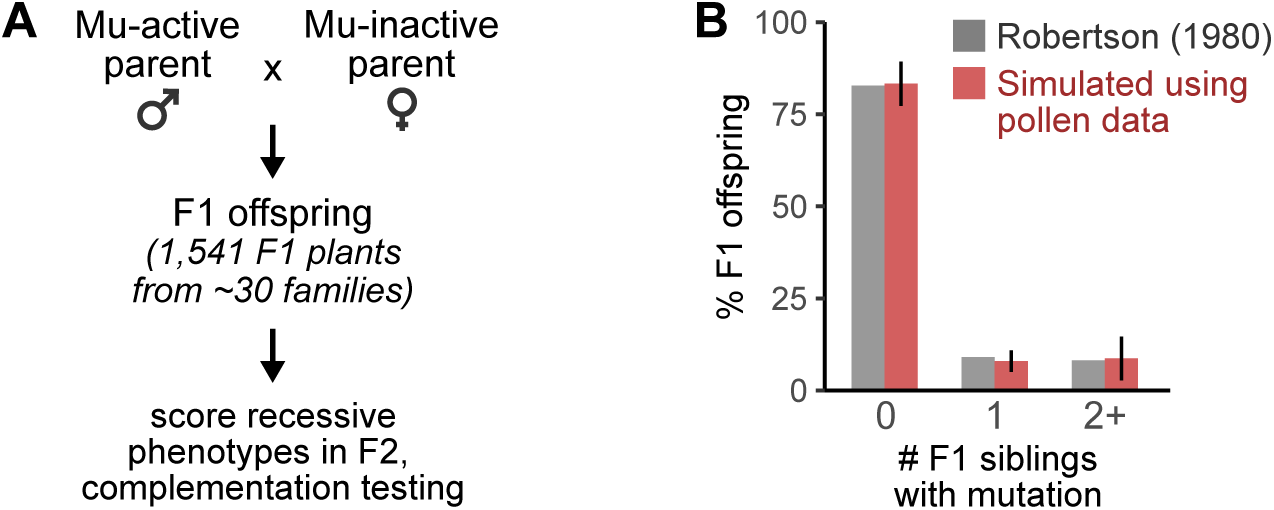
Pollen allele frequencies are consistent with outcross data from classical genetics. (**A**) Experimental design from Robertson (1980). F1 offspring were generated by outcrossing a Mu-active male parent, then the offspring were assessed for appearance of visible mutant phenotypes after self-fertilization. F1 siblings segregating similar mutant phenotypes were subjected to complementation testing to determine if they shared the same (allelic) mutation. (**B**) The experimental design in A was simulated using mutant alleles randomly drawn with probabilities matching the measured pollen allele frequencies. The % of F1 offspring that share mutations with 0, 1, or 2+ siblings were then calculated and compared to Robertson (1980). Error bars, standard error of the mean.

To reconcile these results, we directly compared our data to Robertson (1980). We previously showed that pollen allele frequency data can predict inheritance patterns in the offspring (**Fig. 2**); this approach can also be used to predict more complex experimental designs, such as Robertson’s. We simulated Robertson’s experiment 1,000 times, randomly drawing new mutations at probabilities defined by the bulk pollen data (*SI Appendix*). On average, the simulations predict that 83.3% of F1 offspring would have unique mutations (**Fig. 5B**), in close agreement with the reported value (p = .83, two-tailed bootstrap test). An advantage of the simulated experiments is that it is possible to computationally go ‘back in time’ and see how abundant any given mutation was in the Mu-active parent (**Fig. S14**). For offspring that shared mutations with 2+ siblings, the source mutations had an average allele frequency of 0.13 in pollen (consistent with a mutation at the time the seed was planted^45^); thus, the fact that Robertson observed any such offspring (8.2% of the total) suggests that early Mu activity occurred at an appreciable rate in this experiment.

Here, we can provide an alternative explanation for Robertson’s data: in a dividing cell population, most mutations will be rare simply because there are more cells later in development and therefore more opportunities for a mutation to occur. While there is one chance for a mutation in the zygote, there are two in the following division, then four, and so forth. This effect will lead to increasing numbers of mutations at decreasing allele frequencies, as was observed in all tissues for Mu insertions (**Fig. 3A**) and is even predicted by the Luria-Delbrück distribution (**Fig. 2B, S15**). Rather than evidence for tissue-specific activity, the preponderance of unique mutations in Mu outcross families is better explained by the statistics of cell division.

## DISCUSSION

*De novo* mutations are difficult to identify because they can be extremely rare within a tissue. This has led to an acute depth-vs-breath trade-off^46^, where mutations can either be sequenced to lower depth across the genome or at higher depth for targeted loci. Here, we overcame this technical barrier with a strategic model system – the maize Mu transposon. We show that Mu sequencing can accurately measure the absolute allele frequency of *de novo* TE insertions genome-wide, while achieving a detection limit rivaling the most sensitive of targeted mutation studies^18^.

As a model of mutation, a limitation of our approach is that it is only applicable to TEs; however, several findings are likely to be representative of other mutation classes. First, there were a large number (>100,000) of *de novo* Mu insertions per sample. If single-base substitutions could be sequenced to the same depth, a similar number of events might be expected. It has been estimated that every gene is mutated multiple times in an organism the size of maize or humans^9^, a prediction consistent with data from deep sequencing single-genes^18^. The number of *de novo* Mu insertions per cell was similar to the germline single-base substitution rate in maize^35^ and far below the number of single-base substitutions per somatic cell in animals^34^. Thus, Mu simply provides a glimpse into the scope of genetic mosaicism for an organism with a cell population measured in the trillions.

Second, most *de novo* mutations were present at low allele frequencies. A strong trend towards low frequency mutation is expected from the statistics of cell division, as there are exponentially more cells later in development and thus more chances for mutations to occur. While individually rare, these mutations can collectively add up to important effects and may contribute to aging, cancer, and evolution. Finally, tissues varied not only in the number of mutations per cell, but also in how widespread the mutations were. When considering the rise and spread of *de novo* mutations, it will be important to recognize that multicellular organisms are large, complex populations with extensive heterogeneity.

Our results provide greater resolution into Mu activity across maize tissues. Mu insertions have been observed in somatic tissues such as leaf^29^, but quantitative data on their number and abundance were not available. We found that Mu insertions occurred continuously throughout development in both somatic and germinal tissue. This is in contrast to Mu excisions, as there is a clear bias against early excision activity^40^ and a >1,000-fold range in excision rates between endosperm^40^ and pollen^26^ (vs. 4-fold maximum range for *de novo* insertions). This provides further evidence that Mu insertions can be decoupled from excision outcomes, perhaps due to tissue-specific differences in the use of DNA repair to restore a Mu element after a transient excision event^26^.

What might drive the tissue-specific variation in Mu allele frequencies? Differences in transposon activity may contribute, but are not the only explanation. For instance, spatial biases in cell division rates have a profound impact on mutant allele frequencies^9,11–13^, and so differences in tissue development may contribute to the patterns we observed. Selection for and against specific mutations has been observed in healthy human tissues^16^, and might similarly impact the persistence or spread of *de novo* Mu insertions. Future work can dissect the relative contribution of tissue-specific transposon activity, cell division patterns, selection, and other processes on the ultimate abundance of *de novo* mutations within and across the plant.

## ACKNOWLEDGMENTS

We thank Jonathan Gent, Robert Erdmann, Bob Schmitz, and Chris McFarland for invaluable discussions and critical reading of the manuscript. We thank Grant Freeman for initial testing of the MuSeq2 protocol. We thank the Duke University Sequencing and Genomics Technologies Core for sequencing services. Funding was provided by NIH grant R35GM151237 to B.N.

## AUTHOR CONTRIBUTIONS

Conceptualization, B.N.; Methodology, J.S. and B.N.; Visualization, B.N.; Software, J.S. and B.N.; Formal Analysis, M.H. and B.N.; Investigation, J.S.; Writing – Original Draft, B.N.; Writing – Review & Editing, J.S., M.H., and B.N.

## MATERIALS AND METHODS

### Sample preparation and MuSeq2 libraries

Mu-active and inactive lines were descended from Maize Co-op stocks 919J and 910I (respectively). DNA was extracted using a modified CTAB protocol (endosperm, pollen) or the Qiagen Dneasy Plant Mini kit (leaf, root), and then sheared with a Covaris E220 Sonicator. A MuSeq2-specific adapter (**Table S2**) was ligated onto the DNA with the NEBNext Ultra II DNA Library Preparation Kit, and Mu-containing fragments were then selectively amplified with nested PCR using primers that target a conserved region at the edge of Mu elements (**Table S3**). More detailed methods are available in ***SI Appendix***.

### Data mapping and analysis

Reads were mapped to the W22 genome^47^ and converted to molecular counts using a UMI added during adapter ligation. Paternal insertions were then identified as insertion sites detected at >1000 counts per million in both endosperm and another matched tissue for a given plant. To convert to allele frequencies, the number of UMIs at each insertion site was divided by the mean UMIs for the paternal insertions and then multiplied by 0.33 (for endosperm) and 0.5 (other tissues). Detailed analysis methods are described in ***SI Appendix***.

### Data and code availability

Sequencing data are available at NCBI GEO under accessions GSE279993 and GSE296286. Code for mapping and analysis can be found at github.com/jts34805/Quantitative-and-sensitive-sequencing-of-somatic-mutations-induced-by-a-maize-transposon/.

