## Supplementary material for "Quantitative and sensitive sequencing of somatic mutations induced by a maize transposon": SI Appendix and Figures

Literature survey on VAF detection limit vs. genomic coverage

For **Fig. S1A**, only papers that reported VAFs in the main text or figures were considered. The selected studies were not meant to be exhaustive, but rather representative of a range of techniques and mutation types. VAF detection limits were as given in each paper; when multiple limits were provided for different sample types (e.g. tissues), the lowest reported value was used. For targeted sequencing studies, genomic coverage was calculated from the reported target size divided by the mappable genome size[48]: 2,864,785,220 bp for human and 119,482,012 bp for *Arabidopsis*. For the purposes of the figure, whole-genome sequencing papers were considered to have 100% genomic coverage.

Two classes of study were not included: First, studies that identified *de novo* mutations from transcript data were excluded (e.g. ref 49). RNA-seq produces very uneven read depths across the genome, and so there is not a well-defined relationship between VAF detection limit and genome coverage – highly expressed genes, which represent a small portion of the transcriptome, dominate the minimum observed VAFs. Single-cell mutation studies were also not included[50]. This was not due to active exclusion, but rather because available single-cell mutation papers largely do not report VAF detection limits. While single-cell methods provide additional cell-type resolution and information about cell-to-cell variation, they do not fundamentally overcome the limitations on sequencing depth and error rates that are also present in bulk experiments.

Plant growth and tissue collection

Mu-active plants were maintained by continual outcrossing of Mu-active pollen onto Mu inactive ears, using the bz1-Mum9 anthocyanin reporter to confirm Mu activity. The Mu-inactive maintainer (female) parents were descended from maize Co-op stock 910I, and carry the sh1-bb1981 and bz1-m4::Ds alleles. Mu-active seeds were descended from maize Co-op stock 919J, which carries a mutable bz1-Mum9 allele. Both stocks were originally ordered from the Stock Center in January 2010 by Jonathan Gent. Continual outcrossing of Mu active lines onto Mu inactive ears was required to maintain Mu activity. Mu active kernels were phenotypically identified by the speckling pattern that occurs when Mu somatically excises from the bz1-Mum9 allele.

For tissue collection, kernels were chipped using a razor blade and the resulting endosperm samples were stored in 2 mL tubes and frozen. To minimize sample cross-contamination, the surface that kernels were chipped on was wiped with a 10% bleach solution and razor blades were only used once. Chipped kernels were then planted in vermiculite (Therm-O-Rock Vermiculite 3A-HORT Medium). After the second seedling leaf was fully emerged (V2 stage; 10-13 days after planting), plants were removed from vermiculite. Roots were rinsed thoroughly in water to remove any vermiculite, and the bottommost ~1 inch of primary root (up to and including the root tip) was collected into 2 mL tubes and frozen. The topmost half of the first leaf (~3/4 inch) was also harvested and collected into 2 mL tubes and frozen. Seedlings were then transplanted to soil in the Botany Greenhouses in Athens, Georgia, where they were grown in sunlight supplemented with LED fluorescent lights (Medic Grow 550W Slim Power 2) until maturity. At maturity, pollen was collected into 2 mL tubes in the morning from the plants at first pollen shed (9-10 am) and frozen.

DNA isolation

*Leaf DNA isolation:* Leaf tissue was disrupted in a 2 mL tube using liquid nitrogen and a pestle (Agilent cat. no. PES-15-B-SI). Once disrupted, DNA was extracted with the Qiagen DNeasy Plant Mini Kit (Qiagen cat. no. 69104) and eluted in two steps, first with 30 ul of elution buffer followed by 25 uL elution buffer. DNA size distributions were evaluated using 5 ul of sample on a 0.8% agarose gel.

*Root DNA isolation:* Root tissue was disrupted with a Qiagen Tissue Lyser II and three to six 3 mm glass beads per sample. Prior to disruption, the sample box was chilled overnight at -80 °C with root samples and 3 mm glass beads inside. Pre-chilled root tissue was then shaken in the Tissue Lyser at max frequency for 5 minutes. Samples were removed from the shaker and agitated using a pestle to dislodge root debris from the tube walls. Shaking was repeated at max frequency for 5 minutes. After this process, root DNA was extracted as described for leaf isolation using the Qiagen DNeasy Plant Mini Kit. DNA size distributions were evaluated on a 0.8% agarose gel.

*Endosperm DNA isolation:* Genomic DNA was isolated from endosperm using a modified CTAB DNA extraction protocol. To prepare CTAB buffer, CTAB stock was made with 3% Cetyltrimethyl ammonium bromide, 1.4 M NaCl, 20 mM EDTA (pH 8.0), 100 mM Tris-Cl (pH 8.0). The day of DNA extractions, 2% w/v polyvinylpyrrolidone (PVP, MW 40 kDa) was dissolved into CTAB stock by heating the solution to 65 °C, and then 800 ul preheated lysis buffer was aliquoted into tubes (one tube per sample to be processed). Then 8 ul proteinase k (ThermoFisher cat. no. EO0491) and 1 ul beta-mercaptoethanol were added to each tube.

Endosperm tissue was disrupted using liquid nitrogen, mortar, and pestle. Disrupted tissue was then incubated at 65º C for 1 h in the preheated lysis buffer. Samples were inverted to mix every 10 minutes during incubation. Following this, samples were spun down at 5000 rcf for 8 mins to pellet tissue debris. Lysate was transferred to a new tube using a metal spatula and combined with 1 volume of a 24:1 chloroform isoamyl alcohol solution. Samples were mixed by inversion for 5 minutes and then centrifuged at 8,000 rcf for 10 minutes. The upper aqueous phase was carefully transferred to a new tube following centrifugation. To precipitate DNA, 0.7X volumes of cold isopropanol was added to each sample and inverted to mix. Samples were incubated at -20 °C for 1 hour. Samples were then centrifuged at 10,000 rcf for 15 minutes. The supernatant was removed and the DNA pellet was washed using 1000 ul of freshly prepared 70% ethanol. Samples were inverted to mix and incubated at room temperature for 5 minutes. Samples were then centrifuged for 5 minutes at 10,000 rcf. The ethanol wash was repeated one more time and DNA pellets were dried until the pellet became translucent. The DNA pellet was resuspended using 55 ul of ultra-pure $H_2O$ (ThermoFisher cat. no. 10977015) and incubated overnight at 4 °C. Size distributions were visualized using 5 ul of purified DNA on a 0.8% agarose gel. The first batch of endosperm samples showed signs of cross-contamination; these data were used to identify paternal insertions but excluded from all other analyses (see 'Sample assessment and quality control'). Prior to processing subsequent endosperm samples, the mortar, pestle, and metal spatula were incubated for 5 min in 10% bleach solution and then thoroughly rinsed with water; this additional washing step removed the cross-contamination.

*Pollen DNA isolation:* Pollen was disrupted using a Qiagen Tissue Lyser II as described for root. During disruption, pollen debris would stick to the lid of the tubes and so a pestle was used to scrape off the debris into the tube. After disruption, 800 ul of preheated CTAB lysis buffer (prepared as described for endosperm) was added to each sample. A pestle was again used to scrape off any material from the tube lid back into the tube as well as break up any pellet that had formed in the bottom of the tube. This step ensured a homogenous mixture during lysis, which greatly increased DNA quality and quantity. DNA extractions were then performed as described for endosperm, with the additional of third ethanol wash after DNA precipitation. The first batch of pollen samples had much lower sequencing depth compared to the other tissues (lower UMIs / sample). For subsequent samples, pollen DNA was purified an additional time with a Monarch DNA and PCR cleanup kit (New England Biolabs cat. no. T1030S); the DNA cleanup kit was performed after CTAB extraction and DNA shearing, prior to the end repair step in MuSeq2.

MuSeq2 adapter preparation

MuSeq2 adapter oligos (**Table S2**) were ordered from Integrated DNA Technologies, suspended to 100 uM in TE. The general adapter structure is as follows:

5'-[phos]rrrrrrrrrrUGTGACTGGAGTTCAGACGTGTGCTCTTCCGATCTNNNNNNNNbbbbbbbbbbT-3'

The 10 nt sequences labeled as strings of 'r' and 'b' are reverse complements of each other, allowing the adapter to form a hairpin with a 3' T overhang. These sequences vary by adapter (**Table S2**), providing a sample-specific barcode during adapter ligation. The series of 'N's is the 8 nt Unique Molecular Identifier (UMI). A uracil (U) near the 5' end makes it possible to cut the hairpin after adapter ligation. Read 2 begins at the UMI and continues through the sample barcode and subsequent genome sequence.

To prepare MuSeq2 adapters and anneal the hairpin, 7.5 ul of adapter oligo (100 uM) was diluted with 25 ul Duplex Buffer (Integrated DNA Technologies cat. no. 11-05-01-03) and 17.5 ul H2O. Diluted adapters were then placed in a thermocycler and incubated at 95ºC for 2 minutes followed by a .1º C ramp down in 1-second intervals for 700 cycles, reaching a final temperature of 10ºC. Adapters were then stored at -80ºC.

MuSeq2 library preparation

DNA samples were sheared using a Covaris E220 Evolution instrument in 50 uL of water. Shearing settings were optimized for each tissue to shear to a mean of 1000 bp. All tissues used settings of 2% Duty Factor with 200 cycles per burst. For pollen, the peak incident power was 140 and time was 50 seconds; Endosperm: 100 Peak Incident Power and 30 seconds; Leaf: 70 Peak Incident Power and 30 seconds. Root: 100 Peak Incident Power and 20 seconds. Concentrations and size distributions for sheared DNA was measured using an Agilent 4200 TapeStation with a D5000 screentape (Agilent cat. no. 5067-5589).

Sheared DNA (200-1000 ng / sample) was end-repaired using the NEBNext Ultra II DNA Library Preparation Kit (New England Biolabs cat. no. E7370L) according to manufacturer instructions, except that all reaction volumes were cut in half. MuSeq2 adapters were then ligated to the DNA using the same kit (NEBNext Ultra II) with half reaction volumes; a separate adapter was used for each sample, providing up to 48 sample-specific barcodes during the initial ligation step. After ligation, 1.5 uL USER enzyme (New England Biolabs cat. no. M5505S) and 2 uL Exonuclease 1 (New England Biolabs cat. no. M0293S) were added to each sample and the reaction was incubated at 37 for 15 min then 80 °C for 15 min. This step linearizes the hairpin adapters by cleaving at a uracil base, and the addition of Exonuclease 1 degrades residual unligated adapter to minimize carryover in subsequent PCR. Samples were then purified with Ampure XP Beads (Beckman Coulter cat. No A63880) using a bead:sample ratio of 0.8X. After bead purification, libraries were resuspended in 5 ul ultra-pure H2O.

Adpter ligated libraries were processed through 3 rounds of PCR to selectively amplify Mu-containing fragments and complete the Illumina adapter sequences (PCR primer sequences in **Table S3**). For the first PCR, 5 ul sample was mixed with 6 ul NEBNext Ultra II Q5 Master Mix (New England Biolabs cat. no. M0544S), 0.5 ul TIR6 primer (4.8 uM stock concentration; 0.2 uM final), and 0.5 ul UDz_i7 primer (4.8 uM stock concentration; 0.2 uM final). Reactions were pipetted to mix and incubated at 98 °C for 30 s, then 14 cycles of 98 °C for 10 s, 65 °C for 30 s, and 72°C for 30 s, followed by 72 °C for 2 min. To remove excess primers, 0.5 ul Exonuclease I was added and the tube was incubated at 37 °C for 15 min then 80 °C for 15 min.

For PCR2, an additional 4 ul of Q5 master mix was added along with 0.4 ul of Museq2_NestedTIR primer (10 uM stock), 0.4 ul of P7 primer (10 uM stock), and 2.7 ul of ultra-pure H2O. Reactions were pipetted to mix and incubated at 98 °C for 30 s, then 6 cycles of 98 °C for 10 s, 59 °C for 30 s, and 72°C for 30 s, followed by 72

°C for 2 min. To remove excess primers, 0.5 ul Exonuclease I was added and the tube was incubated at 37 °C for 15 min then 80 °C for 15 min.

For PCR3, 5 ul of PCR2 product was mixed with 25 ul Q5 master mix, 19.5 ul ultra-pure H2O, and 5.5 ul xGen indexed primer pairs (Integrated DNA Technologies xGen UDI Primers Plate 1, cat. no. 10005922). A distinct primer pair was added to each sample to allow for multiplexing. Reactions were pipetted to mix and incubated at 98 °C for 30 s, then 5-15 cycles of 98 °C for 10 s, 59 °C for 30 s, and 72°C for 30 s, followed by 72 °C for 2 min. The number of PCR cycles varied by sample and was determined using qPCR as follows: prior to PCR3, 5 ul of the prepared PCR reaction was withdrawn and mixed with 0.5 ul of a 1:1000 dilution of SYBR Green I DNA Gel Stain (Thermo Fisher cat. no. S7563) in water. The reaction aliquot with SYBR was then run on a BioRad CFX96 Real Time PCR Thermal Cycler for 25 cycles using the reaction conditions listed above. The cycle number for each PCR reaction was chosen to be within 30-80% of the plateau height, and the remaining PCR mix was run using the selected cycle number.

After PCR, libraries were cleaned up and size selected using magnetic beads. For cleanup, 50 ul Ampure XP beads were added to each PCR sample (1X ratio) and then the DNA was purified according to manufacturer instructions and eluted in 40 ul ultra-pure H2O. Size selection was then performed using SPRIselect beads (Beckman Coultier cat. no. B23317) with 0.6/0.8 bead ratios according to manufacturer instructions. DNA was eluted in a final volume of 20 ul and the size distribution and concentration was measured using an Agilent 4200 TapeStation with a D5000 screentape. Libraries were pooled to 15 nM such that each individual library was equally represented. Paired-end 150 bp sequencing was performed at the Duke University Sequencing and Genomics Technologies Core on an Illumina NovaSeq X Plus instrument with 20% PhiX spike-in.

<u>Mu insertion mapping and quantification</u>

For MuSeq2 libraries, read 1 contains the last 29 bp of the Mu transposon followed by genomic DNA sequence. Read 1 was first pre-processed by removing the first 23 bp, which contain transposon sequence matching the PCR primer, and moving bp 24-29 (the 'validation sequence') to the read header using Fastp v0.23.4[51] with all filters disabled. Read 2 contains the adapter ligated fragment with an 8 bp UMI, 11 bp sample-specific barcode, and genomic sequence. Read 2 was pre-processed using Fastp to move both the 8 nt UMI and 11 nt sample-specific barcode to the read header. During this step, adapter sequences were also trimmed and fragments with under 40 bp in read 2 were removed. Paired-end reads were then mapped to the W22 V2 genome[47] using Bowtie2 v2.5.4[52], with both mixed and discordant mapping disabled (--no-mixed --no-discordant). The UMI-tools v1.1.6[53] 'group' function was used to identify reads sharing the same UMI; to account for sequencing errors in the UMIs, the default 'directional-adjacency' method from the UMI-tools package was used (described in ref. 53).

Next, fragments were filtered using a custom R script to remove low quality mapping and PCR duplicates. Fragments were excluded if they had a mapping quality score <10 or a validation sequence that did not match 'TRTCTC' (the sequence at the edge of the Mu transposon). They were further excluded if they did not have an exact match to the sample-specific barcode added during adapter ligation. To remove PCR duplicates, fragments mapping to the same position (both read 1 and 2) with the same UMI were merged. The libraries in this study were intentionally over-sequenced to increase the amount of error-correcting from molecular counting, with a mean of 5.2 sequenced fragments per molecule (UMI). Each molecule was required to have a minimum support of at least 1/5 the average number of reads for a given sample; for the median library, this means that each molecule (UMI) was sequenced with a minimum of two reads.

Most Mu elements contain an intact Terminal Inverted Repeat (TIR) at both ends of the transposon, and so can be sequenced out of each direction. To connect molecules mapping to the left or right border of the same Mu element, the following steps were taken: First, for Mu elements present in the reference genome (N = 20; all historical insertions), the left and right borders were defined based on the genome sequence. For other Mu elements, the left and right borders were connected by expecting a 9 bp target site duplication (TSD) to be generated during Mu insertion; this would result in both ends of the transposon mapping 8 bp apart in reverse orientation (for a 9 bp TSD, there is an 8 bp distance between positions 1 and 9). To allow for discrepancies in TSD length, we searched for cases where a left and right border were within 15 bp of the expected TSD, but where both borders had over 50-fold more molecule counts than the corresponding border 8 bp away. In such cases, the two borders with higher counts were considered to come from the same element. Deviations from the 9 bp TSD were rare, with only 273 such instances identified compared to more than 3 million with the 'ideal' 9 bp TSD. For Mu insertion sites supported by at least 10 transposon-spanning molecules, 92% were sequenced out of both directions and 8% were only supported by one TIR.

The total number of molecule counts mapping to either the left or right transposon border were then added to provide a single estimate for each element. A subset of elements were not sequenced effectively out of both directions, which could result in under-counting as there was only one border available for sequencing instead of the usual two. To adjust for this effect, we identified any elements where there was a greater than 2-fold difference in molecule counts between the left and right border after adding a pseudocount of 500. For these elements, the number of molecules was estimated as 2 times the greater of the left or right border counts. This process affected 422 elements (0.00013%). Finally, 18 elements were 'blacklisted' and removed from analysis (**Table S4**) because they were identified at moderate abundance (between 10-1000 counts per million) in over half of all samples or half of the Mu-inactive controls; many of the blacklisted sites were ancestral Mu elements with diverged sequences and would not be expected to amplify efficiently during MuSeq.

#### Estimating variant allele frequencies from Mu count data

To convert Mu insertion counts to variant allele frequencies (VAF), the data for each sample was first scaled to counts per million (CPM). Paternal insertions were identified as insertion sites with ≥1000 CPM in both endosperm and at least one matched sporophytic tissue (leaf, root, or pollen), excluding the 29 historical insertions (**Table S1**). CPM data were then normalized to VAF by dividing each sample by the mean CPM of the paternal insertions and multiplying by 1/2 (for leaf, root, and pollen) or 1/3 (for endosperm); the difference in normalization factor for endosperm is because the endosperm is triploid with a 2:1 maternal:paternal ratio, and so paternal DNA makes up 1/3 of the DNA in this tissue. The random error for normalization is estimated to be 6.2% (standard error of the mean for the paternal insertion sites).

#### Identifying transmitted *de novo* insertions from two-generation Mu families

Four families were created by crossing a Mu-inactive female parent with a Mu-active male. MuSeq2 libraries were prepared from endosperm of the male parent and both endosperm and leaf of 7-8 offspring as described above. After mapping and quantification, inherited insertion sites were identified as insertion sites with ≥500 CPM in leaf/endosperm of an offspring and ≥200 CPM in the matched leaf/endosperm (e.g. an insertion with 500 CPM in leaf and 200 CPM in the matched endosperm would qualify). Paternal genotype calls were made based on having ≥200 CPM in the paternal endosperm and inheritance in at least one offspring. The reason that a lower CPM threshold was used to genotype insertions in the two-generation families (compared to the other samples, as described in the "Estimating variant allele frequencies" section, above) was that these samples were sequenced to lower depth and the reduced thresholds avoided a few false negatives in the samples with the least sequencing depth. A matrix of genotype calls is available at GEO accession GSE296286. The strategy used to call de novo insertions transmitted from the Mu-active male parent is shown in **Fig. S7**.

Evaluating the sensitivity of MuSeq2

The detection limit for each sample was calculated as the VAF for a Mu insertion supported by a single transposon-spanning molecule, and ranged between $1.5 \times 10^{-5}$ to $3.5 \times 10^{-4}$ (median = $6.0 \times 10^{-5}$).

To estimate the % of Mu insertions captured during library prep, we compared the amount of input DNA to the number of transposon-spanning molecules sequence per paternal insertion (**Table S5**). The input DNA amount was converted into the number of genome equivalents using a conversion factor of 407.37 genome equivalents per ng total DNA; this factor was calculated based on a haploid genome size of $2.4 \times 10^9$ bp, an average molecular weight per bp of 651.98, and assumes all DNA is nuclear (e.g. the contribution of plastid and mitochondrial DNA is negligible). The number of genome equivalents was then converted into the expected number of times a paternally inherited transposon insertion would be recovered, if each transposon insertion contributed a single final molecule. On average, we estimate that 14.0%, 8.3%, 16.5%, and 16.7% of transposon insertions were sequenced in leaf, pollen, root, and endosperm (respectively), not accounting for loss during DNA purification. The lower % yield for pollen is likely explained because an extra column purification was performed after measuring the DNA concentration, while for every other tissue the DNA was directly input into library prep.

*Yield per pollen grain:* Other than yield during library preparation itself, loss can occur during DNA purification or because of incomplete sample loading (most libraries were prepared from a portion of the total isolated DNA). To estimate the total yield for pollen: each pollen sample was collected into a 2 mL tube and filled to the 100 uL mark. By weighing the mass of pollen in several samples collected this way, we estimate average mass of pollen per sample was 120 mg. To determine the number of pollen grains per mg, we diluted pollen samples in defined volumes of water and then counted the number of pollen grains in a 5 uL aliquot. From this, we estimate there are 1750 pollen grains / mg, or ~210,000 pollen grains per sample (1750 pollen grains / mg x 120 mg / sample). If every Mu insertion were sequenced once, then we would expect a pollen sample to result in 315,000 transposon-spanning molecules per heterozygous paternal insertion (210,000 pollen grains x 3 copies of the genome / pollen grain x 0.5 heterozygous insertions / genome copy); we observed an average of 4052 molecules per paternal insertion in pollen (**Table S5**), and so ~1 in 78 insertions were sequenced.

*Estimated yield at each step of MuSeq2:* With 100% yield after DNA purification, one pollen sample would thus produce 1550 ng total DNA (210,000 pollen grains x 3 genome equivalents / pollen grain / 407.37 genome equivalents / ng). After CTAB purification, we obtained an average of 630 ng DNA, or 41% yield. Libraries were then prepared from 45% of the total DNA; if higher total % yield were desired, then the entire DNA fraction could be used by simply scaling up volumes and using more enzyme. A second column purification was then performed (only for pollen), and we estimate a yield of 53% for the column purification by assuming that the reduced number of molecules sequenced for pollen compared to all other samples is explained by the extra column purification step (see above). Finally, there was a ~7.5% yield for converting DNA to sequencable insertions (15% yield per Mu insertion divided by 2 ends per Mu element = 7.5% yield per transposon border). One major source of loss during library preparation is the presence of sheared fragments where the transposon end is not the correct distance from the fragment border. DNA was sheared to 1000 bp and the final libraries were size selected to 250-500 bp (100-350 bp genomic DNA insert + 150 bp adapter sequence). If the transposon end were randomly distributed relative the the sheared DNA end, we would expect it to fall within 100-350 bp of the end 25% of the time, and so a 4-fold loss in yield is explainable simply by random shearing. As a result, the 7.5% yield during library prep is 25% yield from shearing size and 30% yield from incomplete conversion during adapter ligation, cleanup, and PCR. In summary:

~210,000 pollen grains → 41% yield, CTAB DNA purification → 45% of DNA used for library prep →

53% yield, column cleanup → 15% yield per insertion (7.5% yield per transposon end) during library prep

Contribution of pre-meiotic insertions to mutations transmitted through pollen

A transposon insertion during or after meiosis could occur in a maximum of 6 molecules (if the insertion were before meiotic S-phase). Given that only 1 in 78 insertions were sequenced, the probability of sequencing a meiotic or post-meiotic insertion more than once is nearly zero ($p = 0.003$; Poisson distribution with lambda = 6/78). Therefore, any insertions sequenced more than once occurred prior to meiosis. Out of ~22.5 insertions per pollen grain (Fig. 2, 4), an average of 19 (85%) were supported by insertions sequenced more than once ($\Sigma vaf_i$ for all insertions supported by ≥2 molecules). The remaining 15% can be attributed to a combination of meiotic, post-meiotic, and pre-meiotic insertions. At the detection threshold (78 molecules), over half of the remaining divisions are pre-meiotic (3-4 mitotic divisions + meiosis + 2 pollen divisions = ~78 molecules after replication). If the Mu transposition rate was equal throughout these late divisions, this would imply that slightly over half of the remaining 15% of insertions are pre-meiotic and the other half are post-meiotic. There are several uncertainties in estimating how to divide up this final 15% of insertions, including the potential for changing Mu activity over time and the fact that not all pollen grains were collected (the 210,000 pollen grain samples were maybe 25% of the total pollen shed in a given day). Conservatively, we estimate a minimum of 1/3 of the remaining insertions occur during the late pre-meiotic divisions and so the contribution of pre-meiotic insertions is >90% (best estimate, ~95% of insertions are pre-meiotic).

Sample assessment and quality control

In total, 46 Mu-active samples were collected and sequenced for this study. All samples were used when identifying inherited insertions (paternal and historical insertion sites), but 17 were excluded from further analysis because of concerns with library quality: 6 samples were excluded because they did not meet a minimum VAF detection threshold of $10^{-3}$. This set included the first 5 pollen samples, which had consistently low molecule counts, and one endosperm sample. Subsequent pollen libraries incorporated an additional round of DNA purification (see 'DNA isolation' section, above) and resulted in much better sequencing depth. Second, the first 11 endosperm libraries were excluded because they showed evidence of cross-contamination; in these libraries, paternal insertions from one library consistently showed unusually high abundance in the others. For subsequent endosperm libraries, all non-disposable items used for tissue disruption (mortar, pestle, metal spatula) were subjected to a more stringent washing protocol that included soaking in 10% bleach for 5 min (see 'DNA isolation' section, above); this additional cleaning step resolved the cross-contamination issue. These sample exclusion criteria were set prior to analyzing the data further.

Interpreting the allele frequency distribution of *de novo* Mu insertions

To estimate the mean allele frequency distribution for each tissue (e.g. **Fig. 4A**), the cumulative number of *de novo* Mu insertion sites at or above a given VAF was first calculated for the individual samples. The single-sample allele frequency distributions were then log-transformed and interpolated at 200 evenly spaced points between $\log_{10}(10^{-5})$ and $\log_{10}(1)$ using the R function *approx* (R version 4.3.0). The mean and 95% confidence interval (CI95) for each tissue was then calculated by bootstrapping with 2000 bootstrap replicates. As the sequencing depth varied between samples, the mean was reported down the minimum VAF covered by at least 75% of samples in a tissue. Power-law fits to the allele frequency distributions were performed using the R *lm* function after $\log_{10}$ transformation.

Simulating the Robertson (1980) experiment from pollen allele frequency data

Robertson (1980) performed a series of outcrosses between Mu-active plants (F0) and Mu-inactive donors. The F1 progeny were then evaluated to determine if they segregated new mutations and whether any of the mutations were shared with siblings. In total, 1541 F1 offspring were tested, of which there were 171 mutant plants (11.1%) carrying an estimated 154 distinct mutations.

To simulate the results of one Mu outcross from Robertson (1980), a MuSeq pollen sample was first randomly selected. This sample represents a single Mu-active F0 plant from Robertson's study. Mutations (Mu insertion sites) were then randomly drawn based on the measured pollen allele frequencies. For instance, a mutation with a VAF of 0.1 was drawn with a 10% chance of occurring in each F1 offspring. After simulating 50 such F1 offspring (roughly the average number of offspring evaluated per outcross in ref. 27), the number of times a mutation occurred 1, 2, or >3 times among the offspring was recorded. Robertson's entire study had ~30 such outcrosses, for a total of 1541 F1 plants. Thus, to simulate a full iteration of Robertson's study, 30 simulated outcross experiments were performed using 30 different pollen samples (randomly sampled with replacement) and the totals were added together.

To estimate confidence intervals, it is important that the simulated study reflects the variation expected under the conditions of Robertson (1980). From the pollen data, an average of 32,038 mutations were recovered for each simulation, far more than the 154 mutations recovered by Robertson (1980). This discrepancy is explained because Robertson tracked mutations with visible seedling phenotypes, which would represent only a small portion of the total. To better match the counting noise during Robertson (1980), the simulated mutations were downsampled so that an average of 154 were recovered per simulation. This downsampling makes the simulation-to-simulation variation better matched to Robertson (1980), but does not affect the mean estimates: 83.1% of mutations were found to be unique prior to downsampling, compared to 83.3% after downsampling.

Simulating the Robertson (1980) experiment assuming a Luria-Delbrück process

The Robertson (1980) experiments was simulated assuming a Luria-Delbrück process (constant mutation rate under exponential cell division) for Fig. S15. To simulate the results of one Mu outcross from Robertson (1980), plant growth under mutation was simulated as a series of 16 mitotic divisions, followed by meiosis and two pollen mitotic divisions. The choice of 16 mitotic divisions was because this is the minimum number of divisions required to obtain the population size of a typical pollen sample in this study (~250,000 pollen grains corresponds to 2^16 pollen mother cells after the mitotic divisions and then 4 pollen grains per meiosis). Prior to each cell division, a random number of mutation was simulated according to a poison process with lambda (mutation rate) equal to 2 * (171/1541) / 19 = 0.0117; this mutation rate results in a matched number of mutant F1 offspring as observed by Robertson, where the factor of 2 is because of meiotic reduction (there will be twice as many mutations in the diploid pollen mother cells as the number found in offspring), 171/1541 is the proportion of F1 mutant plants identified by Robertson (171 mutant F1 out of 1541 F1 plants), and the factor of 19 accounts for the total number of simulated cell divisions (16 mitotic divisions + meiosis + 2 pollen divisions). For meiosis, a single round of mutation was followed by two divisions and then mutations were retained with a probability of 0.5 (this simulates meiotic reduction where there is a 50% chance of any given mutation appearing in a meiotic product). Pollen mitoses were simulated with the same mutation rate but without an ensuing cell division as only the lineage leading to the fertilizing sperm cell will contribute to the offspring. After simulating cell division and mutation, 50 'cells' were randomly drawn and the number of mutations shared by 0, 1, or 2+ sibling was calculated. This was repeated 30 times to simulate a single round of the Robertson (1980) experiment, and then the whole experiment was simulated 1000 times to determine the mean and standard error of the mean.

### SI FIGURES

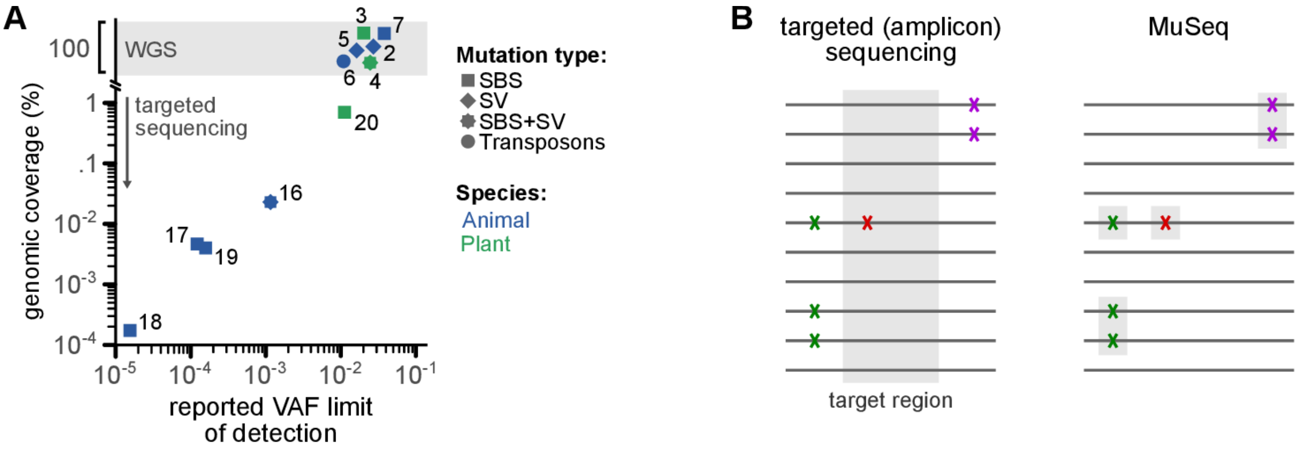


**Figure S1. Sequencing depth limitations make it difficult to assess rare *de novo* mutations**

(**A**) Relationship between VAF detection limit vs genomic coverage for selected studies. Numbers reflect the citation number in the main text references. Here, '100% genomic coverage' implies there was not intentional selection for a subset of the genome; in practice, this means the 'mappable genome' and excludes regions that are repetitive or otherwise difficult to amplify or sequence. VAF, variant allele frequency; SBS, single-base substitution; SV, structural variant.

(**B**) Comparison between targeted (amplicon) sequencing and MuSeq. While both approaches limit the sequencing to a portion of the genome, they do so in different ways. In these cartoons, 10 example DNA sequences are illustrated as dark gray lines; there are three mutations at different abundances, colored as green, red, and purple 'X's. For targeted sequencing, the 'X's could represent any class of mutation (SBS, SV, transposon); for Museq, these must be Mu transposon insertions. The region targeted by each technique is highlighted in gray.

Targeted sequencing selects a predefined set of genome loci to sequence deeply. Both wild type and mutant alleles are sequenced and any mutations outside of the target region are missed. MuSeq, in contrast, sequences transposon insertion sites throughout the genome, and reduces sequencing depth by avoiding the wild-type (transposon-free) alleles. In this hypothetical example, targeted sequencing would require at least 10 reads but only capture a single mutation; MuSeq would require fewer reads yet would capture all 3 mutations. While MuSeq is limited to transposons (Mu in this case), the opportunity is that it enables orders of magnitude greater sensitivity and dynamic range than is possible for other classes of mutation.

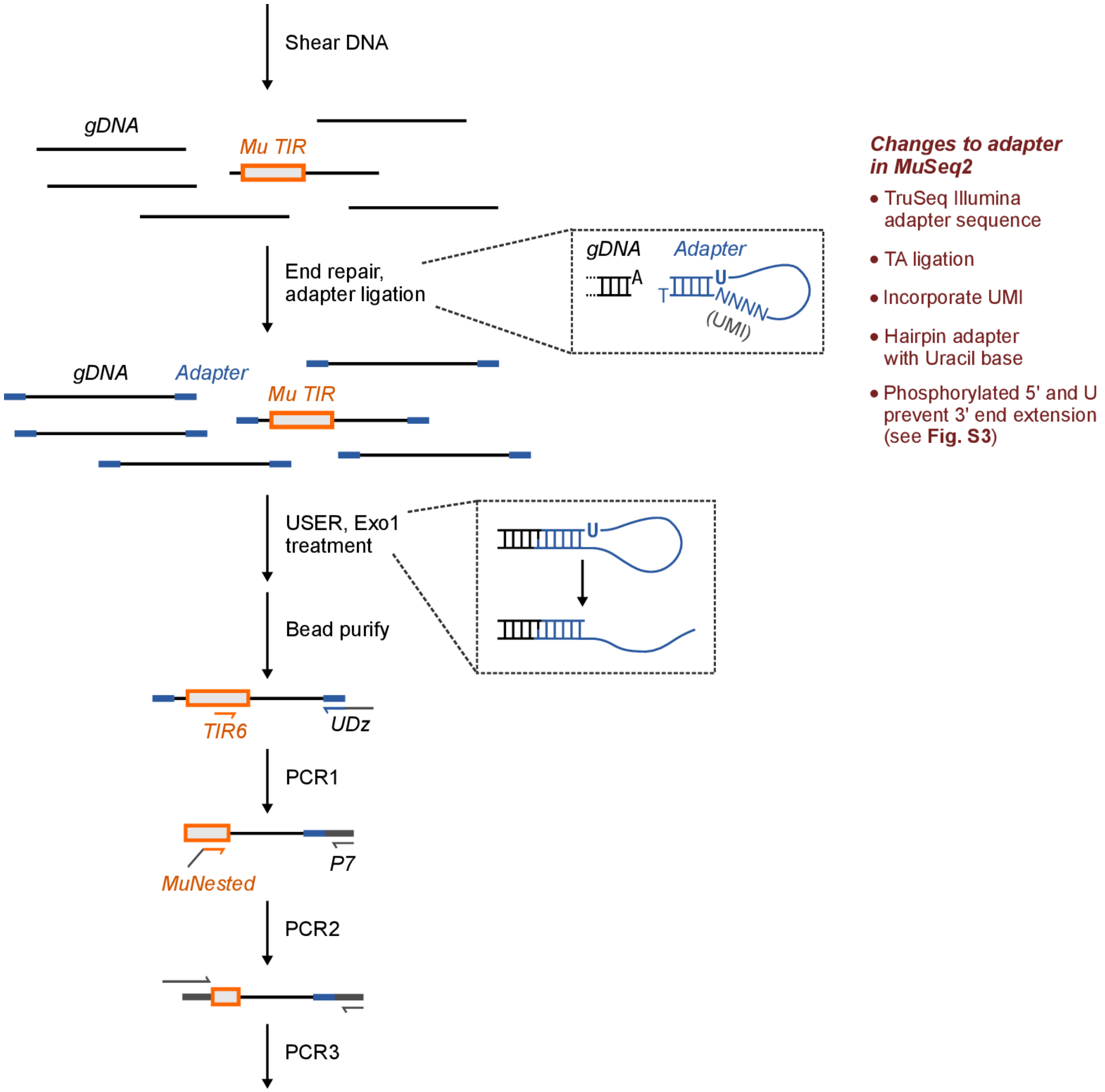


**Figure S2. Overview of the MuSeq2 protocol**

MuSeq2, similar to MuSeq, uses an adapter ligation followed by a series of nested PCR reactions to specifically amplify fragments spanning the transposon genome junction. Several changes to the adapter were made in MuSeq2, including incorporating a Unique Molecular Identifier (UMI) for molecular counting and updating to modern Illumina adapter sequences. USER treatment cleaves the adapter at the Uracil, making the ends compatible with the downstream PCR reactions.

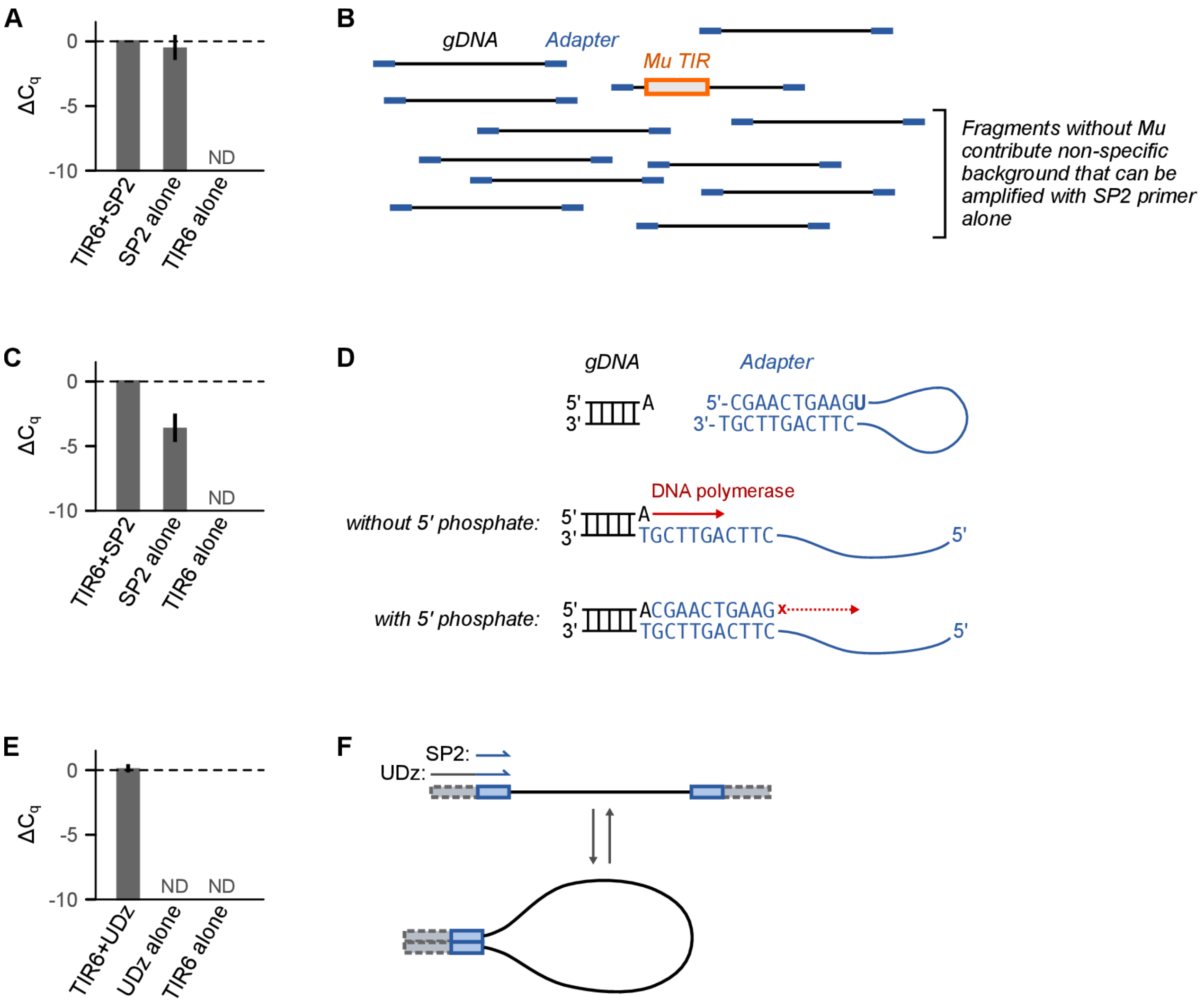


**Figure S3. Reducing non-specific amplification through changes to adapter structure and suppression PCR**

(**A**) Quantitative PCR for libraries prepared using an adapter design similar to the original MuSeq: the adapter was unphosphorylated and the reverse primer (SP2) matches the adapter-ligated sequence only. TIR6 is a Mu-specific primer; the SP2 primer matches the Illumina sequence ligated onto sheared genomic DNA. Three independent DNA samples were sheared and ligated, then the ligated DNA was split for qPCR with different primer combinations. The amplification is not specific, as SP2 alone amplifies similarly to when the Mu-specific primer was included. Cq was normalized to TIR6+SP2; in this experiment, the average number of cycles at ΔCq = 0 was 16.4.

(**B**) The reason for non-specific amplification with the SP2 primer is that the majority of DNA fragments from sheared genomic DNA do not contain a Mu element. Fragments without Mu are estimated to outnumber Mu-containing fragments by more than 10,000 fold. The background fragments will have adapter DNA on both sides, forming a potential priming site for PCR with the SP2 primer. The adapter structure limits background amplification in part because it has a 5’ overhang and does not initially have the sequence needed for primer binding (the primer binds to the reverse complement of the overhang; this was also true in the original MuSeq); however, the 5’ overhang can be copied by DNA polymerase at the start of PCR. This is likely an inefficient process, but given the excess of fragments without Mu it still contributed meaningful background (panel A).

(**C**,**D**) One modification to reduce non-specific background in MuSeq2 was to replace the unphosphorylated oligo with a phosphorylated one. With a 5’ phosphate on the adapter, both adapter strands can be ligated to the sheared genomic DNA. After a Uracil on the adapter is cleaved to release the hairpin, it leaves a 3’ phosphate overhang. By ligating the adapter on both strands, there is no free 3’ hydroxyl available – blocking extension by DNA polymerase. Panel **C** shows the same experiment as panel **A**, except using a phosphorylated adapter. Cq was normalized to TIR6+SP2; in this experiment, the average number of cycles at ΔCq = 0 was 16.2.

(**E**,**F**) A second modification in MuSeq2 was to use a longer primer, UDz, in place of SP2 during the first PCR. Fragments with adapter sequence on both sides do not amplify as efficiently because they have self-complementary ends and can form a hairpin (suppression PCR). The UDz primer adds the entire Illumina adapter sequence during PCR. Because this primer makes the self-complementary region longer, it favors hairpin formation and increases the amount of suppression PCR for non-specific fragments. Panel **E** shows qPCR using the same adapter-ligated samples as in panel **C**, except that PCR was performed with the UDz primer. Cq was normalized to TIR6+SP2 from panel **C** and so the ΔCq values are directly comparable between these panels. There was no decrease in specific amplification when switching to the UDz primer, but non-specific amplification was completely suppressed.

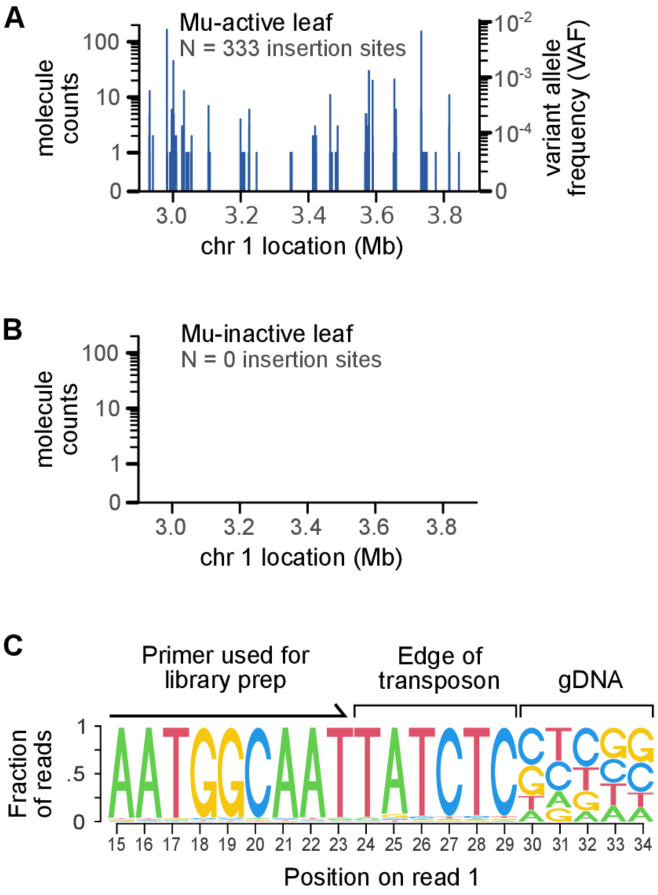


**Figure S4. Specificity of MuSeq2**

(**A**) A representative 1 Mb region showing all insertion sites identified in a single Mu-active leaf sample. In total, 333 insertion sites were found in this region, covering a range of molecule abundances (UMI counts). The y-axis on the left shows the normalized molecule counts while the axis on the right is normalized to variant allele frequencies (VAF). Molecule counts and VAF are directly proportional to each other; for this sample, the normalization was based on an average of 5860 +/- 240 molecule counts per heterozygous (VAF = 0.5) paternal insertion.

(**B**) The same region as in **A**, but for a Mu-inactive leaf sample. No insertion sites were observed in this region.

(**C**) Sequence composition for a portion of read 1, which covers the transposon-genome junction. The last 6 bp of the transposon were not included in any primer used during library prep, and provides independent validation that the sequencing is specific to Mutator. The ‘validation sequence’ matches the known transposon sequence TATCTC for the vast majority of reads.

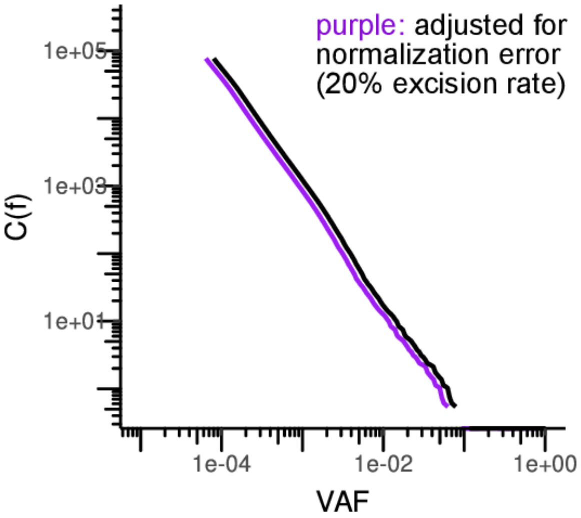


**Figure S5. Mu excision rates have a negligible impact on normalization**

Allele frequencies were normalized using the paternal insertions, which were assumed to be at their original abundance (VAF = ½ in most tissues, 1/3 in endosperm). In the presence of excisions, the true allele frequency of paternal insertions would be less, resulting in systematic error during normalization. We estimate the endosperm excision rate in our line is ~10%, based on the proportion of endosperm surface that has reverted to purple (this line carries a mutable bz1-Mum9 reporter allele that allows for purple pigment expression after excision). To be conservative, we used double this rate – 20% – and calculated the effect this would have on the measured allele frequencies. This figure shows the allele frequency distribution for a representative endosperm sample before (black line) and after (purple line) adjusting for normalization error due to a 20% excision rate.  Even with double the observed excision rate, normalization error has a minimal impact on the allele frequency spectrum. This is because a change on the order of 20% is small when the measured frequencies vary by many orders of magnitude (log-scale). As a result, we did not consider excision further in our analyses; all main text results were not adjusted for excision (e.g. the black line above).

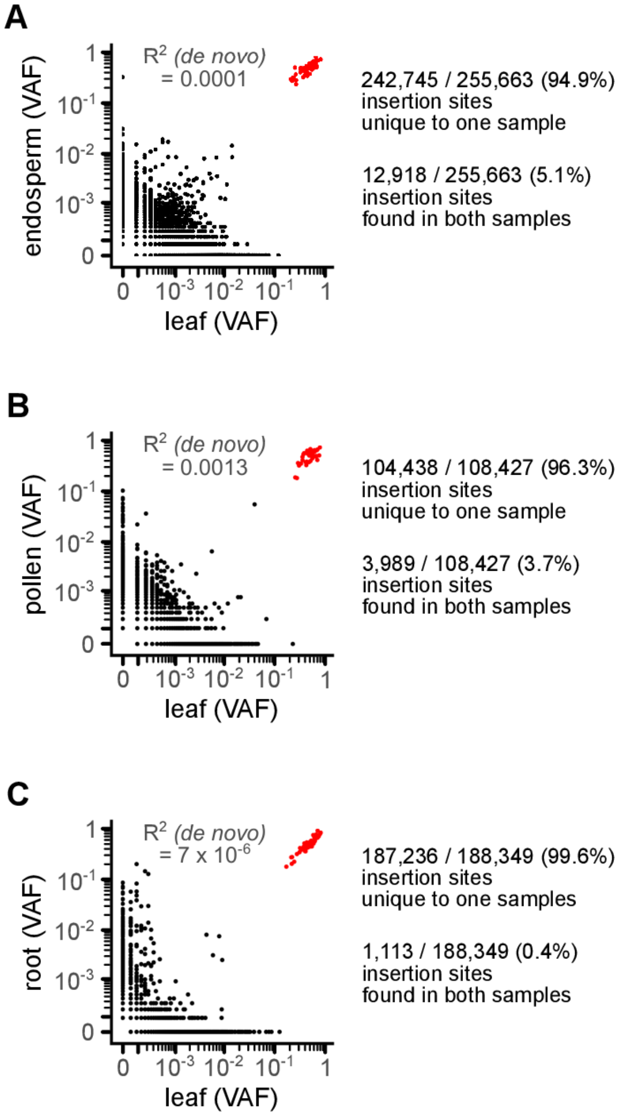


**Figure S6. Comparison between insertion sites identified in matched tissues from the same plant**

Panel (A) is the same as main text Fig. 1C, except that the number of insertion sites unique to one sample (either leaf or endosperm) vs shared between both are quantified. Panels (B) and (C) are similar, except they show a comparison between leaf and pollen (B) or root (C) from the same plant.

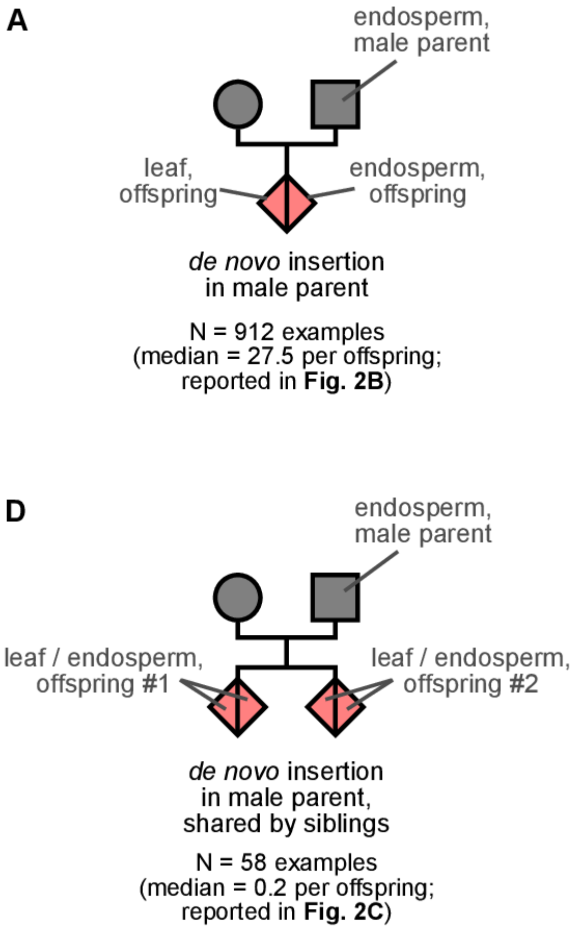


**Figure S7. Transmission of *de novo* Mu insertions into the offspring.**

Mu insertions were sequenced in the endosperm of Mu-active male parents and both endosperm and leaf of the offspring (N = 7-8 offspring per family, 4 families in total). The female in each of these crosses was Mu-inactive, and so new Mu insertions can be attributed to the male parent.

(**A**) *De novo* Mu insertions that occurred in the male parent were identified based on their absence in paternal endosperm and presence in both endosperm and leaf of the offspring. There was a median of 27.8 ± 6.7 *de novo* Mu insertions per generation.

(**B**) Mu insertions inherited by the offspring but present in the paternal endosperm could be attributed to prior generations (grandparents and beyond) and were not *de novo* in the male parent.

(**D,E**) *De novo* insertions from the male parent were most often transmitted uniquely to a single offspring (**E**) and only occasionally were shared by siblings (**D**). The low rate of *de novo* insertions shared by siblings was reported by Robertson (1980) and also explored in **Fig. 5**.

(**C,F**) To assess the contribution of technical errors to the genotyping calls, we looked for cases where an insertion occurred in two samples that cannot be explained by genetics. For instance, there were 914 cases where an insertion was present in the leaf of an offspring and the endosperm of its parent. This situation can be explained if the parent transmitted the insertion to the offspring, and so we would expect the matched offspring endosperm to also have the same insertion. This was seen in every case (**B**) and we never observed the biologically impossible situation where a matched endosperm did not have the insertion (**C**). Similarly, we observed 5,760 cases where the leaf and endosperm of one plant had an insertion not present in a sibling (**E**) but 0 cases where an insertion was present in the leaf of one plant and the endosperm from the sibling (**F**).

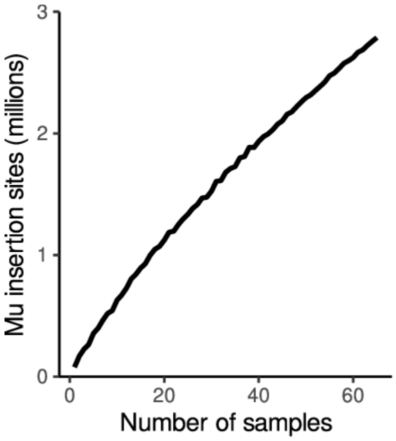


**Figure S8. The number of Mu insertion sites has not reached saturation**

Random subsets of samples were drawn and then the total number of genomic insertion sites was calculated. The total number of insertion sites has not reached saturation under the conditions of this study.

**A**

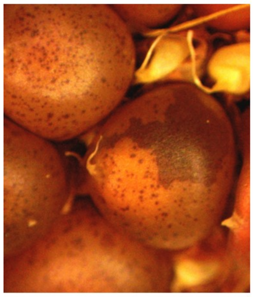

**B**

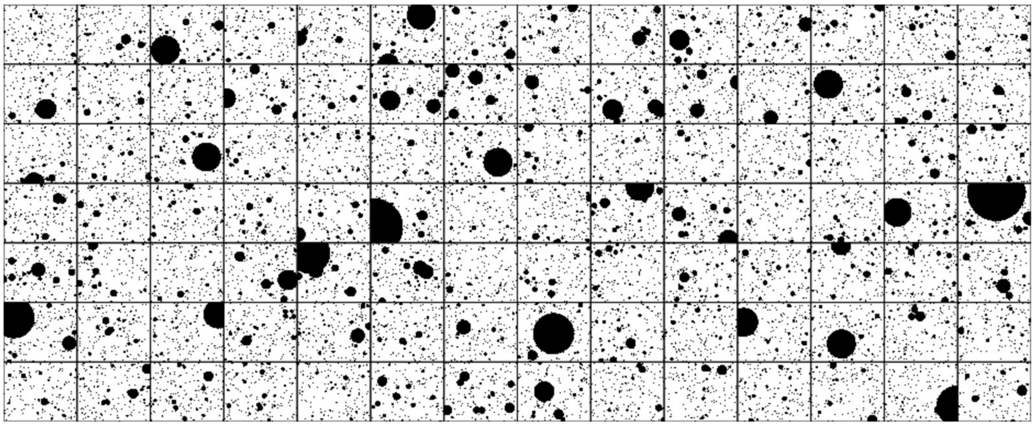

**Figure S9. Mu insertions and excisions behave differently in endosperm**

(**A**) Example sector sizes in Mu-active kernels with the bz1-Mum9 reporter. The kernel on the top left with many small spots is representative. The very large sector on the kernel in the middle is exceedingly rare; we observed 7 out of 1844 kernels with sectors making up at least 5% of the kernel area. Prior quantitative data on excision spot size found even fewer large sectors, with 0 sectors at a frequency under $2^8$ (VAF ~ $10^{-3}$) out of 2000 kernels (Levy and Walbot, 1990).

(**B**) A simulated ear with sectors drawn to represent Mu insertions. In this simulation, we assumed all divisions happen within a 2D plane, which may be approximately true for aleurone (the outer cell layer of endosperm where the visible pigment is produced). Spot size was defined by randomly drawing *de novo* insertions based on the measured allele frequencies in endosperm, requiring an average of 10% of the surface to be covered by sectors (10% surface coverage matches the rate measured for Mu excisions from reporter alleles; Levy and Walbot 1990). The frequency of large spots in this diagram is dramatically higher than what has been observed for endosperm excision sectors; in the simulation, 45% of ‘seeds’ had at least one sector covering more than 5% of the surface area, compared to 7/1844 (0.4%) for excisions (panel A).

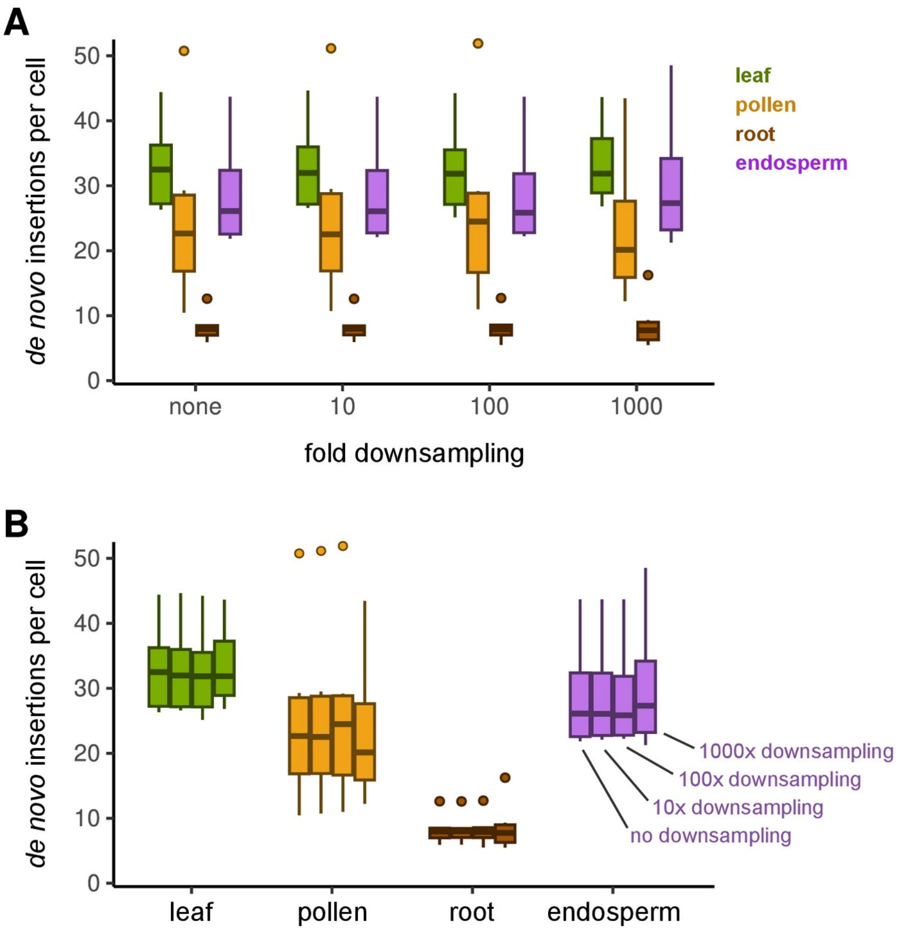


**Figure S10. Estimates for the number of Mu insertions per cell are robust to sequencing depth, relevant to Figs. 4B.**

Samples were downsampled by randomly removing molecules and then repeating the analysis in Fig. 4B. Panels **A** and **B** are identical, except that panel **B** groups all samples from the same tissue together. The estimated number of *de novo* insertions per cell was unaffected by downsampling as much as 1000-fold.

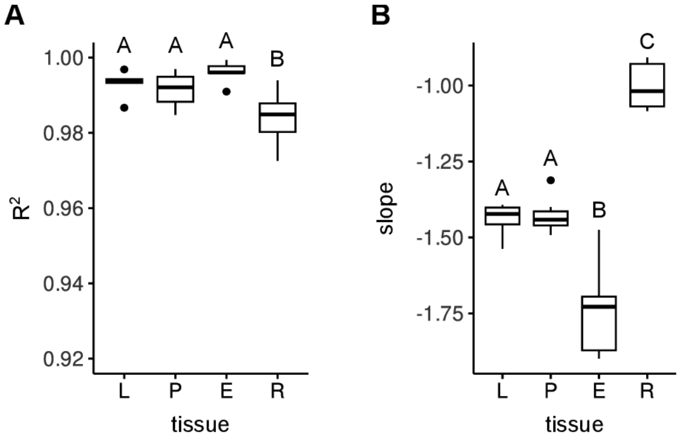


**Figure S11. Power-law fits to the allele frequency spectra from various tissues**

Best linear fit parameters to log-transformed data. Fitting was performed on individual samples and the results plotted as a boxplot separated by tissue. Letters indicate statistical significance: groups not sharing a letter have a significantly different mean ($p \leq 0.05$; Tukey's honest significant difference test). N = 6 samples for leaf and root; N = 9 samples for endosperm and pollen.

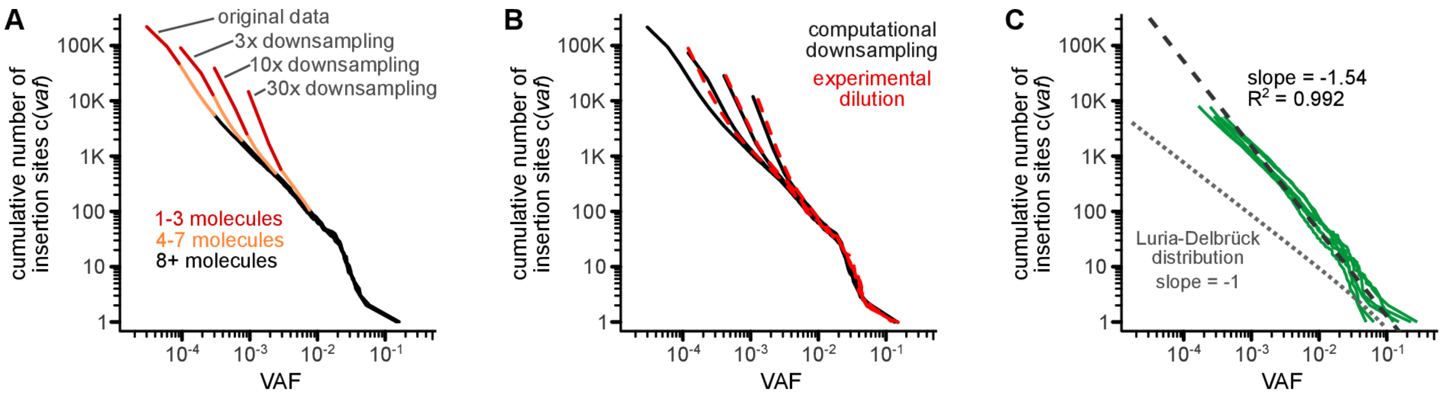


**Figure S12. Evaluating the impact of counting error and sampling statistics on the measured allele frequency distributions.**

(A) The data for a representative leaf sample was down-sampled 3, 10, or 30-fold by randomly drawing transposon-spanning molecules from the total. The down-sampled data were then normalized and the allele frequency distribution was plotted, as in the main text. The portion of the curve supported by 1-3 molecules are shown in red, 4-7 molecules in orange, and 8+ molecules in black. There is an upward bias in the estimated number of insertion sites for the portion of the curve supported by few molecules. The reason is that many insertions can be present below the detection limit, and while these are individually at low abundance and unlikely to be sequenced, they are collectively numerous (e.g. an insertion present at 1/10 the detection limit has a low chance of being sequenced, but there are so many such insertions that some of them will be sequenced once or twice). Beyond 8+ molecules, this effect is negligible (black curves).

(B) DNA from the leaf sample in (A) was experimentally diluted with Mu-inactive DNA, so that the contribution from the Mu-active leaf sample was 100%, 30%, 10%, or 3% of the total. MuSeq2 libraries were then prepared from these diluted samples and sequenced. The allele frequency distribution for the diluted samples (dotted red lines) closely match the distribution predicted by computationally downsampling (black lines). This shows that sampling statistics, such as approximated by random downsampling, capture the major source of bias in these curves.

(C) To assess whether our conclusions were robust to the upward bias in the number of insertions at low allele frequencies, we repeated the analysis in Fig. 3B after removing the data points supported by fewer than 8 molecules. The $R^2$ and slope of the best fit line were similar whether using the full data (1+ molecules; Fig. 3B) or truncated data (8+ molecules; this panel).

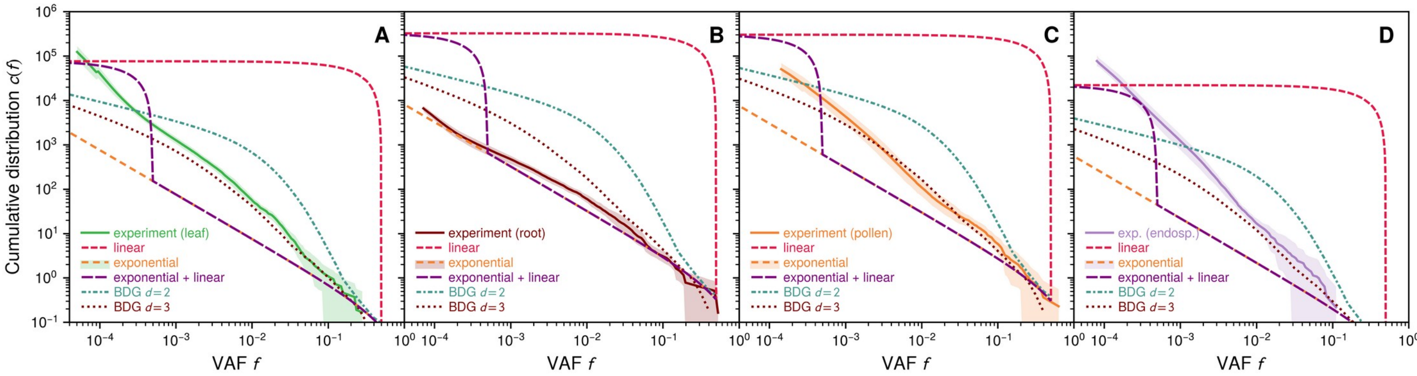


**Figure S13. Fit of experimental leaf data to various models of mutation accumulation**

Details of the theoretical models are described in the SI Text. All models assume no cell death, a constant mutation probability $\mu = 0.076$ over time (chosen to agree with the experimental curve at the largest frequency), and a final cell population size of $N_{cell} = 10^6$. Linear = linear growth, where after each cell division only one daughter cell is capable of further cell division. Exponential = exponential growth, where both daughters are capable of division (Luria-Delbrück model). Exponential + linear = 10 generations of exponential growth, with the remaining generations linear. BDG = boundary-driven growth simulations based on the Eden model, which were carried out on both 2D and 3D square lattices.

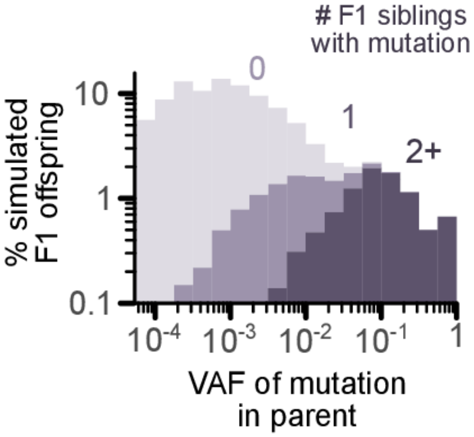


**Figure S14. During simulations of Robertson (1980), F1 offspring that share the same mutation as their sibling are most often derived from pollen insertions at high allele frequencies.**

For the simulations of Robertson (1980) in **Fig. 5B**, we recorded the VAF for any Mu insertion transmitted to the simulated F1 offspring. In this histogram, the bars are color coded based on whether the pollen mutation was inherited by 0, 1, or 2+ F1 siblings. Simulated F1 offspring were derived from Mu insertions at a wide range of allele frequencies, suggesting these occurred throughout development. For mutations present in larger clusters of F1 offspring (2+ siblings with the same mutation), the average VAF in the parent was 0.13; this corresponds 1 insertion in every 3.8 diploid pollen progenitors, roughly the number of meristematic cells in the seed that ultimately form the maize tassel (the male flower; ref: Poethig, Coe, and Johri, 1986). VAF, variant allele frequency.

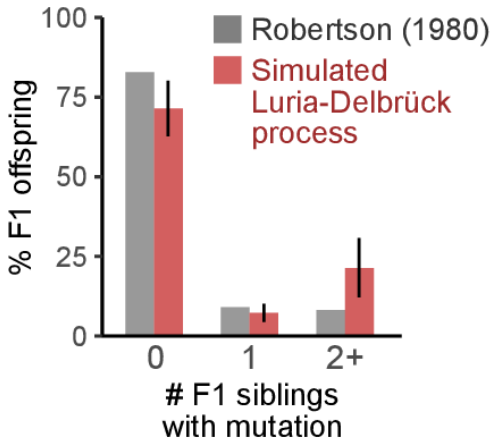


**Figure S15. Comparison between Robertson (1980) to simulations with exponential cell division and a constant mutation rate (a Luria-Delbrück process).**

The Robertson (1980) experiment was simulated assuming a constant mutation rate during 16 mitotic divisions, followed by meiosis and two pollen divisions. Sixteen mitotic divisions was selected because this is the minimum required to achieve the population size of a pollen sample in our study (~250,000 pollen grains). Simulated pollen grains were then drawn according the experimental design of Robertson (1980), and the frequency that an F1 offspring shared a given mutation with 0, 1, or 2+ siblings was quantified. The Luria-Delbrück process predicts fewer F1 offspring with unique mutations compared to Robertson (1980), but this difference is not statistically significant ($p = 0.198$; two-tailed test based on the result of 1000 simulations).
